# TidyMass2: Advancing LC-MS Untargeted Metabolomics Through Metabolite Origin Inference and Metabolic Feature-based Functional Module Analysis

**DOI:** 10.1101/2025.05.09.652992

**Authors:** Xiao Wang, Yijiang Liu, Chao Jiang, Zinuo Huang, Hong Yan, Sunny Wong, Caroline H. Johnson, Jingxiang Zhang, Yifei Ge, Feifan Zhang, Renfu Lai, Peng Gao, Xuebin Zhang, Xiaotao Shen

## Abstract

Untargeted metabolomics provides a direct window into biochemical activities but faces critical challenges in determining metabolite origins and interpreting unannotated metabolic features. Here, we present TidyMass2, an enhanced computational framework for Liquid Chromatography-Mass Spectrometry (LC-MS) untargeted metabolomics that addresses these limitations. TidyMass2 introduces three major innovations compared its predecessor, TidyMass: (1) a comprehensive metabolite origin inference capability that traces metabolites to human, microbial, dietary, pharmaceutical, and environmental sources through integration of 11 metabolite databases containing 532,488 metabolites with source information; (2) a metabolic feature-based functional module analysis approach that bypasses the annotation bottleneck by leveraging metabolic network topology to extract biological insights from unannotated metabolic features; and (3) an intuitive graphical interface that makes advanced metabolomics analyses accessible to researchers without programming expertise. Applied to longitudinal urine metabolomics data from human pregnancy, TidyMass2 identified diverse metabolites originating from human, microbiome, and environment, and uncovered 27 dysregulated metabolic modules. It increased the proportion of biologically interpretable metabolic features from 5.8% to 58.8%, revealing coordinated changes in steroid hormone biosynthesis, carbohydrate metabolism, and amino acid processing. By expanding biological interpretation beyond conventionally MS^2^ spectra-based annotated metabolites, TidyMass2 enables more comprehensive metabolic phenotyping while upholding strict open-source principles of reproducibility, traceability, and transparency.

## Introduction

Metabolomics has emerged as a powerful approach in biomedical research, enabling comprehensive profiling of small molecules that serve as direct signatures of biochemical activities compared to genomics, transcriptomics, and proteomics^1–6^. The human metabolome represents a complex intersection of small molecules derived from diverse origins, such as endogenous metabolism, gut microbiota, environmental exposures, dietary intake, and pharmaceuticals, each of which can independently and significantly influence human health^7–12^. Within this intricate landscape, the gut microbiome alone produces thousands of metabolites that can enter circulation to impact systemic health^7,8,13^. Similarly, xenobiotics from the environment, bioactive compounds from foods and botanical sources, and pharmaceutical agents introduce additional layers of metabolic complexity^7–12^. However, the field faces significant challenges in data interpretation and biological contextualization that limit its full potential.

Despite significant advances in LC-MS untargeted metabolomic technologies, identifying the origin of metabolites remains challenging^14,15^. This limitation significantly hinders our ability to interpret metabolomic data in the context of host-microbiome interactions and environmental exposures. Several computational approaches have attempted to address this limitation, including MicrobeMASST^16^ and MetOrigin2^17^. However, MicrobeMASST is designed for specific microbiota-derived metabolite analysis and is restricted by its reliance on MS^2^ spectra matching, leaving the majority of metabolic features without the source information since only 10-15% of metabolic features typically have MS^2^ spectra^18,19^. Meanwhile, MetOrigin2 can only analyze data from a limited set of metabolite databases and lacks integration with comprehensive metabolomic workflows^17^.

Moreover, metabolite annotation poses a significant challenge in LC-MS untargeted metabolomics, despite the development of numerous advanced methods^14,20,21^. Current MS^2^ spectral-based approaches typically annotate only approximately 10%-20% of detected metabolic features^22^, severely limiting our understanding of dysregulated metabolic functions in physiological and pathological situations. Network-based approaches such as Mummichog^23^ and PIUMet^24^ have attempted to address this limitation through alternative annotation strategies. However, both mummichog and PIUMet rely solely on mass-to-charge ratios (*m/z*) for rough metabolite annotation without incorporating retention time (RT) information, a critical parameter that could substantially reduce annotation redundancy and improve accuracy.

We previously developed TidyMass, a comprehensive R-based computational framework for LC-MS untargeted metabolomics that addressed issues of reproducibility, traceability, and transparency through a consistent tidy data structure and modular workflow^25^. TidyMass provided integrated solutions for data processing, data cleaning, statistical analysis, and pathway analysis, but still faced limitations in metabolite origin inference and extracting comprehensive biological insights from unannotated metabolic features. Here, we present TidyMass2, a significant advancement of the original framework with several critical enhancements that address the aforementioned challenges in LC-MS untargeted metabolomics data analysis.

TidyMass2 introduces several major improvements over its predecessor, TidyMass. First, we have expanded functionality through new functions and packages, most notably the *massdatabase* package^26^, which facilitates seamless management and integration of metabolite, reactions, and pathway databases with existing metabolite annotation (*metid*^27^) and pathway analysis (*metpath*) packages in the TidyMass ecosystem. Second, we have built a comprehensive metabolite database (MetOriginDB) containing 532,488 metabolites with biological source information, enabling sophisticated origin inference for metabolites. Third, we have implemented an improved metabolic feature-based functional module analysis, which provides more comprehensive insights into dysregulated metabolic networks and modules without requiring MS^2^ spectra-based metabolite annotation for all dysregulated metabolic features. Finally, we have developed an intuitive Shiny application that operates across major operating systems (Windows, macOS, Linux), making TidyMass2 accessible to researchers without extensive programming experience while maintaining compatibility with both personal computers and server environments.

Additionally, TidyMass2 remains entirely open-source, with all code and databases (metabolites, pathways, and metabolic network) freely available (https://github.com/tidymass; https://www.tidymass.org/databases/), promoting reproducible and transparent metabolomics data analysis within the scientific community. We anticipate that these enhancements will substantially benefit LC-MS untargeted metabolomics research across diverse biomedical applications.

## Results

### Development and Enhancements in TidyMass2

TidyMass2 represents a significant evolution from its predecessor, TidyMass^25^, leveraging its robust foundation while incorporating a suite of substantial improvements designed to overcome existing limitations in the analysis of LC-MS untargeted metabolomics data. These enhancements encompass key areas such as expanded functionality, enhanced usability, and more powerful analytical capabilities, collectively transforming TidyMass2 into a considerably more comprehensive and versatile computational framework for researchers in the field (**Table 1** and **Figure 1a**). Tidymass2 addresses the growing complexities and demands of modern untargeted metabolomics experiments, offering a more streamlined and insightful approach to data analysis and interpretation.

**Figure 1.**
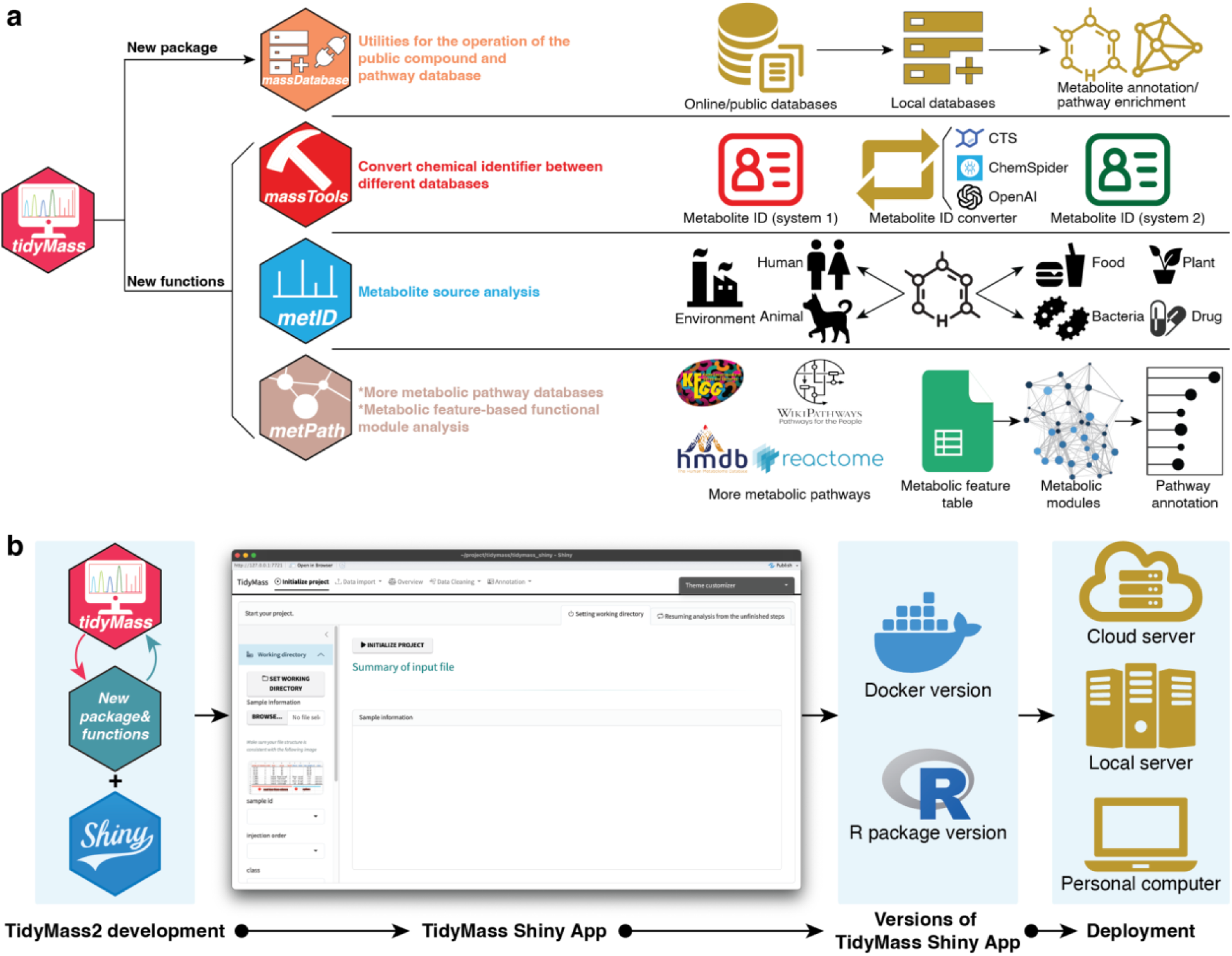
| Development and enhancements in TidyMass2. **a**, Overview of the major improvements in TidyMass2. Key enhancements include the *massdatabase* package for managing metabolomics databases; expanded functionality in masstools for chemical identifier conversion across multiple database systems; metabolite origin inference capabilities in *metid* for tracing metabolite origins; and expanded pathway databases and metabolic feature-based functional module analysis in the *metpath* package. **b,** The *TidyMassShiny* application development and deployment. The left panel illustrates how TidyMass2’s new packages and functions are integrated with Shiny to create the web-based interface. The central panel shows the *TidyMassShiny* user interface for complete data processing and analysis. The right panels demonstrate available deployment options, including the Docker version and R package version that can be implemented on cloud servers, local servers, or personal computers.

**Table 1.** Comparison of key capabilities: TidyMass *vs.* TidyMass2.

| Capability | TidyMass | TidyMass2 |
| --- | --- | --- |
| Metabolite databases with metabolite source information | N.A. | MetOriginDB: 532,488 metabolites with source information (81,461 metabolites with MS<sup>2</sup> spectra) |
| Pathway databases | 2 databases | 4 databases |
| Metabolite origin inference | N.A. | Built-in function in the metabolite annotation pipeline |
| Chemical ID conversion | N.A. | Multi-service system with LLM integration |
| User interface | Command-line only | Command-line and web interface |
| Metabolic feature-based functional module analysis | N.A. | Yes |

#### Comprehensive database management through massdatabase

We have developed *massdatabase*, a new R package that serves as a central hub for online and public metabolite, pathway, and reaction database management^26^. This package enables users to efficiently access, download, and manipulate common online metabolomics databases, including HMDB^28^, KEGG^29^, LipidMaps^30^, DrugBank^31^, FooDB (www.foodb.ca), WikiPathways^32^, and so on (**Figure 1a** and **Supplementary Table 1**). The streamlined integration with existing *metid*^27^ and *metpath* packages creates a seamless workflow from raw data processing to biological interpretation. This integration significantly reduces the technical barriers to comprehensive metabolite annotation and pathway analysis.

#### Enhanced chemical identifier conversion system

Unlike genes and proteins, metabolites lack a universal identifier system in the community^33^. In TidyMass2, we incorporate a chemical identifier conversion system that operates across multiple metabolite ID systems, including HMDB, KEGG, PubChem, ChEBI, CAS, and InChIKey (**Figure 1a**). We have integrated three complementary services to maximize coverage and accuracy: Chemical Translation Service^34^, ChemSpider^35^, and a Large Language Model (LLM) approach based on ChatGPT. This multi-layered approach provides comprehensive chemical identifier conversion, a crucial functionality for cross-database analyses that was previously unavailable in TidyMass.

#### An intuitive graphical interface of TidyMass2

To enhance accessibility for researchers without programming expertise, we developed the *TidyMassShiny* package in TidyMass2, which is a web-based interface that encapsulates all functions and modules available in the command-line version of TidyMass2 (**Figure 1b**). *TidyMassShiny* provides an intuitive graphical interface for the complete LC-MS untargeted metabolomics data processing and analysis workflow, from raw data processing to biological interpretation. Additionally, the Shiny application supports deployment on personal computers, cloud services, and local servers (**Figure 1b** and **Methods**), with detailed documentation for installation and operation across different platforms. This development significantly lowers the technical barriers to utilizing advanced metabolomics data analysis tools.

#### Comprehensive metabolite origin inference

A major advancement in TidyMass2 is the integration of metabolite source information across 11 carefully curated metabolite databases, which is essential for metabolite origin inference (**Methods** and **Figure 1a**). The final comprehensive metabolite database (MetOriginDB) we developed here categorizes metabolites according to their origins, such as human endogenous metabolism, gut microbiome, dietary intakes, pharmaceutical compounds, environmental exposures, and plant-derived substances. Based on the MetOriginDB, we have created a unified reference system that enables researchers to trace the potential origins of metabolites from biological samples. To enhance annotation accuracy, we combined this source information database with both in-house and public MS^2^ spectral libraries. This integration allows TidyMass2 to provide not only metabolite annotation but also origin inference analysis, addressing a critical gap in current metabolomics analysis tools. The ability to trace the metabolite origins offers valuable insights for studies investigating host-microbiome interactions, exposome, and nutritional studies. This part will be elaborated upon in subsequent sections.

#### Expanded pathway databases and metabolic feature-based functional module analysis

Compared to TidyMass, we have expanded the pathway databases within *metpath* to include WikiPathways^32^ and Reactome^36^ pathways, complementing the existing KEGG and SMPDB databases (**Figure 1a** and **Supplementary Table 1**). This expansion provides broader coverage of biological processes and enables more comprehensive pathway enrichment analysis. We calculated the similarities between all the pathways from four different databases and found that combining the four pathways together could significantly increase the number of covered pathways (**Methods** and **Supplementary** Figure 1a,b). Additionally, we leveraged the *massdatabase* package^26^ to download all the reaction pair information from eight established databases (**Supplementary Table 1**). Then we integrated them to build a unified metabolic network (**Methods** and **Figure 1a**). This metabolic network facilitates more analyses, including dysregulated metabolic module identification and targeted enzyme identification. The construction of this metabolic network underpins the metabolic feature-based functional module analysis in TidyMass2, enabling more comprehensive biological interpretation even when MS^2^ spectra-based metabolite annotation is limited. This part will be elaborated upon in subsequent sections.

Collectively, these enhancements make TidyMass2 a more powerful, accessible, and comprehensive tool for LC-MS untargeted metabolomics research. The improvements address key limitations in metabolite origin inference and comprehensive biological interpretation while maintaining the commitment to reproducibility, traceability, and transparency that characterized the original TidyMass framework. Furthermore, in contrast to existing tools utilized for LC-MS untargeted metabolomics^17,23,24,37,38^, TidyMass2 presents several unique advantages, including the provision of customized metabolite/pathway databases, integrated metabolite origin inference, and metabolic feature-based functional analysis (**Supplementary Table 2** and **Methods**).

### Comprehensive Metabolite Origin Inference

Metabolite origin inference addresses fundamental challenges in metabolomics research, particularly for investigating host-microbiome interactions, environmental exposure research, and nutritional interventions^39–41^. Accurately determining whether a metabolite originates from human metabolism, gut microbiota, diet, or environmental exposure provides critical context for biomarker discovery and mechanistic interpretation. For example, differentiating microbial metabolites from host metabolites can reveal specific microbiome functions that could affect host physiology, while identifying environmental contaminants can help assess exposure risks^17^. While several databases containing metabolite source information exist, they exhibit significant limitations in coverage, MS^2^ spectral data integration, compatibility with end-to-end metabolomics workflows, and none of them are open-source^16,17^.

To address these limitations, we have developed a comprehensive metabolite origin inference approach integrated within TidyMass2. We systematically downloaded data from 11 major metabolite databases containing direct or indirect source information, including HMDB^28^, KEGG^29^, DrugBank^31^, ChEBI^42^, FooDB (www.foodb.ca), BiGG models^43^, MiMeDB^44^, Reactome^36^, T3DB^45^, LOTUS^46^, and ModelSEED^47^ (**Figure 2a, Supplementary Table 1**). Each database contributes unique metabolite classes and metabolite source information. For example, HMDB predominantly covers lots of endogenous human metabolites, KEGG provides pathway context across organisms, Drug Bank offers pharmaceutical compound data, and specialized resources like MiMeDB^44^ focus specifically on microbial metabolites (**Methods**).

**Figure 2.**
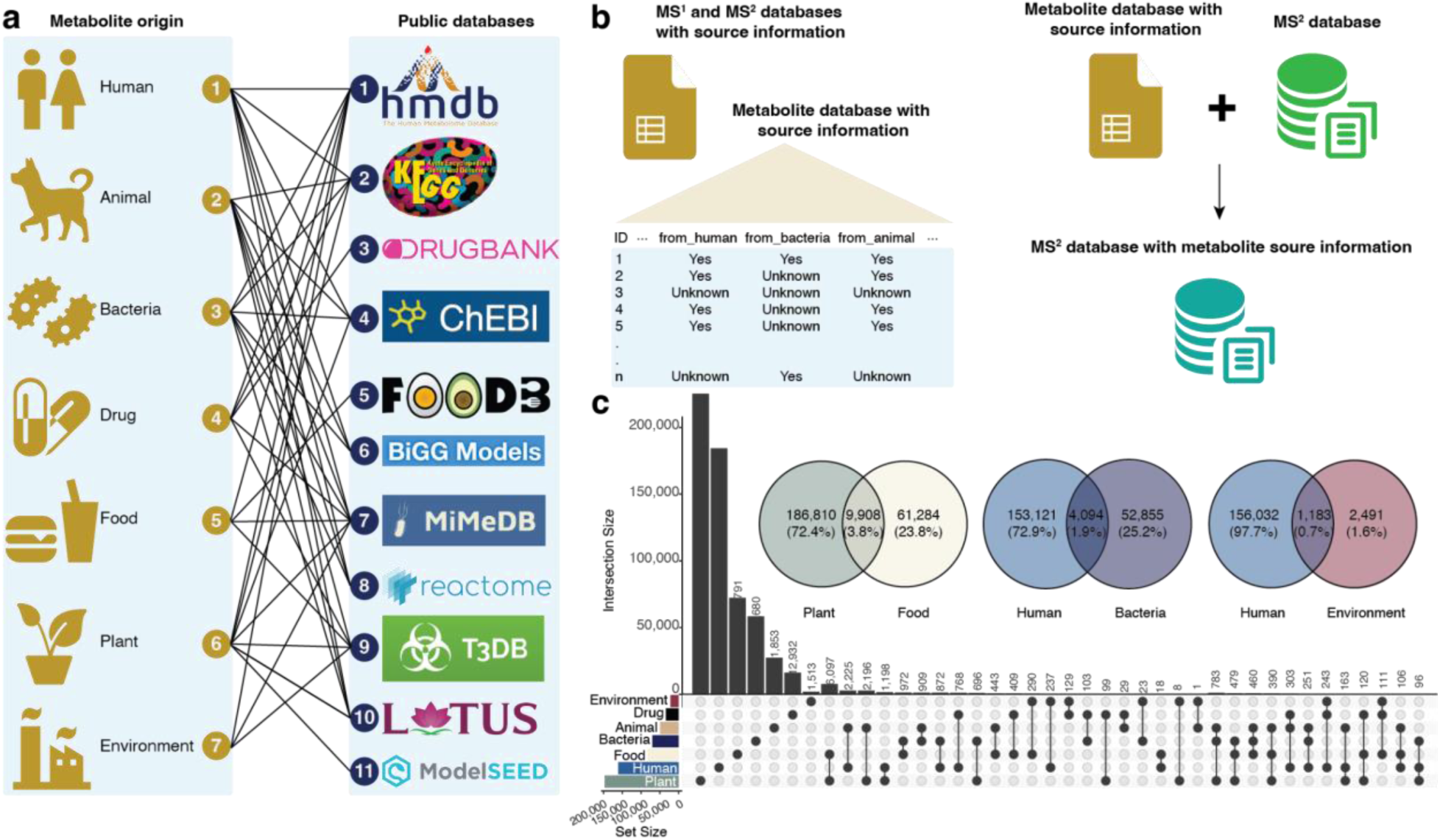
| MetOriginDB: the comprehensive metabolite database with source information. **a**, Source information classification scheme and integration of public metabolite databases. TidyMass2 incorporates data from 11 major databases to enable comprehensive metabolite origin inference across seven source categories. **b,** Framework for integrating metabolite source information with MS^2^ spectral databases. **c,** The overlap between metabolites from different origins.

We categorized all metabolite source information into seven distinct classes: human, animal, bacteria, drug, food, plant, and environment (**Figure 2a**). Moreover, we also documented the precise source information for each metabolite when feasible (**Methods**). For instance, when a metabolite is identified as being of human origin, we will also note the specific derived source (such as blood, urine, placenta, *etc.*) when possible. This classification system enables researchers to precisely track metabolite origins, which is particularly important for distinguishing endogenous compounds from those derived from diet, microbiota, or environmental exposures. After thorough data integration and elimination of redundancies, the final metabolite database contains source information for 532,488 unique metabolites (MetOriginDB), making it the most extensive metabolite database with souce information to our knowledge (**Supplementary** Figure 2a-j) ^17^.

To enhance practical utility for untargeted metabolomics research, we matched the expanded MetOriginDB to in-house and public MS^2^ spectral databases^25^ (**Figure 2b**). This integration creates a seamless connection between metabolite annotation and metabolite origin inference, enabling researchers to perform end-to-end analysis within the TidyMass2 framework. The final MS^2^ spectral database with source information contains 81,461 metabolites with MS^2^ spectra, including approximately 20,156 human metabolites, 15,864 bacterial metabolites, and substantial coverage of other source categories (MetOriginDB_ms2) (**Supplementary** Figure 3a,b).

In MetOriginDB, plant sources contribute the largest number with 196,718 metabolites, followed by human (157,215 metabolites), food (71,192 metabolites), and bacteria (56,949 metabolites) (**Figure 2c**). This distribution demonstrates the diverse origins of metabolites^48–50^. We then explored the overlap between metabolites from different sources (**Figure 2c**). A substantial overlap of 9,908 metabolites was observed between plant and food sources, which was the most significant overlap across all source categories. This is unsurprising given the many varieties of plant-based food^51,52^.

We identified 4,094 metabolites classified as both human and bacterial in origin (**Figure 2c**). This finding quantitatively confirms extensive convergent metabolism between host and microbiome through metabolites^53,54^, highlighting the challenge of distinguishing truly endogenous human metabolites from those produced or modified by gut bacteria^55^. Many bile acids, short-chain fatty acids, and amino acid derivatives exist at this host-microbiome interface, underscoring the concept of “co-metabolism,” where metabolites may be produced through multi-organism metabolic pathways^8^.

Additionally, 1,183 metabolites were found to overlap between human and environmental sources, indicating that environmental exposures may influence the composition and levels of human metabolites. For instance, our database categorizes metabolites/compounds such as urea and acetone as originating from both human and environmental sources. These are typical metabolites produced by the human body, but are also commonly found in the environment^56^.

Through seamless integration with the existing TidyMass2 framework, this comprehensive metabolite origin inference approach maintains the reproducibility, traceability, and transparency that characterized the original platform while substantially extending its analytical power for metabolite origin inference. The MetOriginDB and metabolite origin inference approach in TidyMass2 provides researchers with a powerful tool to contextualize metabolomic findings within the complex interplay of host biology, microbiome activity, diet, and environment.

### Metabolic Feature-Based Functional Module Analysis

Metabolite annotation is still a major bottleneck in LC-MS untargeted metabolomics, with only ∼10–20% of detected features typically annotated via MS^2^ spectral matching^57^. However, the vast majority of unannotated metabolic features (∼80%-90%) still contain valuable biological information and potential biomarkers^58^. This limitation substantially hinders the biological interpretation of LC-MS untargeted metabolomics data and restricts our ability to understand underlying mechanisms^59^.

While accurate mass-based annotation (*m/z* or MS^1^) theoretically enables the annotation of most of the metabolic features, it suffers from high false positive rates, as a single mass can match numerous potential metabolites^60^. To address this fundamental challenge, we developed a metabolic feature-based functional module analysis approach in TidyMass2. This method leverages the inherent network structure of metabolism (metabolic network) to improve annotation confidence and biological insight without requiring comprehensive MS^2^ spectral-based metabolite annotation.

Our approach is based on the principle that metabolism functions as an interconnected network where metabolites are converted to one another through enzymatic reactions. In biological systems experiencing physiological or pathological changes, specific proteins/enzymes are typically up-or down-regulated, causing dysregulation of directly connected metabolites^61^. These perturbations propagate through the metabolic network in a wave-like pattern, resulting in clusters of dysregulated metabolites that form distinct modules rather than random distributions across the metabolic network^62^.

To implement this metabolic network-based approach, we constructed a comprehensive human metabolic network by integrating reaction pairs from 8 public metabolic databases (**Figure 3a and Supplementary Table 1**). After removing the redundant reaction pairs, the final metabolic network comprises 9,630 metabolites connected by 30,196 edges, representing the largest human metabolic network currently available to our knowledge (**Methods and Figure 3b**). Analysis of network topology revealed that most metabolites (72.53%) have connectivity degrees between 1 and 10, consistent with the scale-free properties observed in most biological networks (**Figure 3c**).

**Figure 3.**
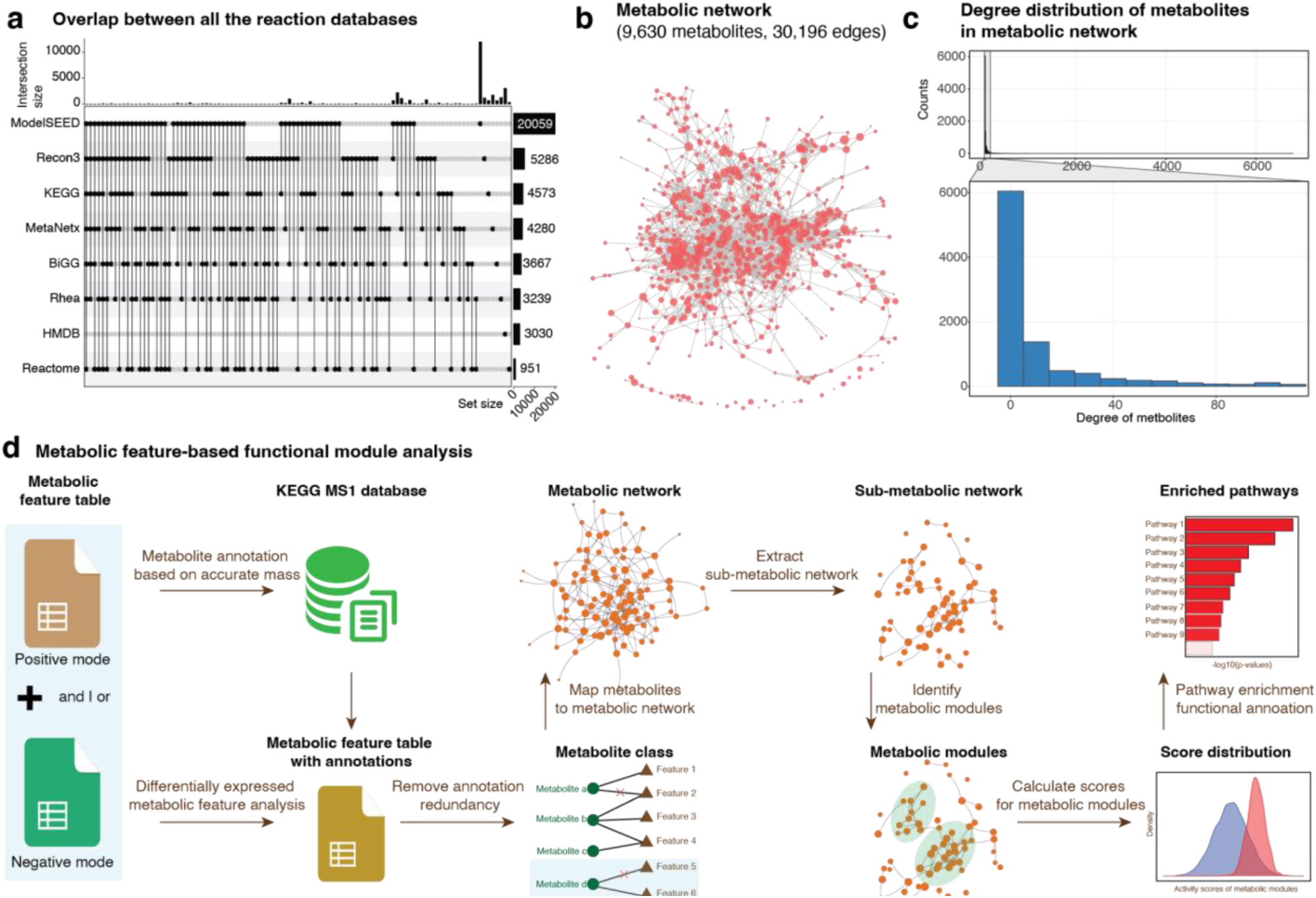
| Metabolic feature-based functional analysis. **a**, The overlap of reaction pairs from 8 databases. **b,** The comprehensive metabolic network was constructed by integrating eight public resources. **c,** Degree distribution of metabolites in the integrated metabolic network. **d,** Workflow of the metabolic feature-based functional module analysis.

Our metabolic feature-based functional module analysis follows a multi-step process (**Figure 3d and Methods**). First, all metabolic features are matched to potential metabolites based on accurate mass (MS^1^). These putative annotations are then grouped into metabolic classes based on chemical annotation and retention time (RT) behavior. A scoring system evaluates annotation confidence, and redundant annotations are filtered according to defined rules (**Methods**). The remaining putative metabolites are mapped onto the comprehensive metabolic network to extract relevant sub-metabolic networks for further analysis.

To identify biologically meaningful modules within this sub-metabolic network, we apply community detection algorithms that partition the network into functionally related metabolic modules. The statistical significance of each identified module is assessed to distinguish true biological annotations from random associations. Metabolic modules showing significant dysregulation are combined to form a dysregulated metabolic network representing the system-level metabolic response. Finally, pathway enrichment analysis is performed for each metabolic module to provide functional context and biological interpretation.

This approach offers several key advantages over pathway analysis using the traditional MS^2^ spectra-based metabolite annotation approaches. Leveraging network topology increases confidence in MS^1^-based metabolite annotations by prioritizing those that form coherent biological modules. It enables biological interpretation even when most individual metabolic features lack MS^2^ spectral-based annotations. Additionally, it provides a comprehensive and systems-level view of metabolic dysregulation that captures the coordinated nature of metabolic responses to biological perturbations. This approach is particularly valuable for mechanistic investigations where complete MS^2^ spectra-based metabolite annotation remains challenging.

### The Diversity of Sources of Urine Metabolites During Human Pregnancy

Pregnancy is accompanied by extensive physiological and metabolic adaptations, many of which can be non-invasively monitored through urine metabolomics^63–65^. Beyond endogenous metabolic changes, emerging evidence suggests that metabolites produced by the vaginal microbiome can influence pregnancy outcomes^66–68^, and that environmental exposures may enter the maternal system and modulate gestational processes^69^.

We selected pregnancy as a case study for TidyMass2 based on several compelling considerations. First, pregnancy is a highly dynamic physiological state characterized by extensive metabolic remodeling across multiple organ systems. Second, the intricate interplay between maternal metabolism, the developing fetus, and the maternal microbiome makes metabolite origin inference especially pertinent. Third, urine provides a non-invasive and information-rich biospecimen, making it an ideal matrix for demonstrating the power of our TidyMass2 framework.

To elucidate the origins of urinary metabolites during pregnancy, we analyzed a longitudinal urine metabolomics dataset using the TidyMass2 framework^70^. This dataset consisted of weekly urine samples collected from a pregnancy cohort across gestation^70^. A total of 875 metabolites were identified through MS^2^ spectral annotation. After excluding those lacking standard identifiers (HMDB, KEGG, or CAS IDs), 662 metabolites retained source classification information and were subjected to further analysis (**Figure 4a and Methods**).

**Figure 4.**
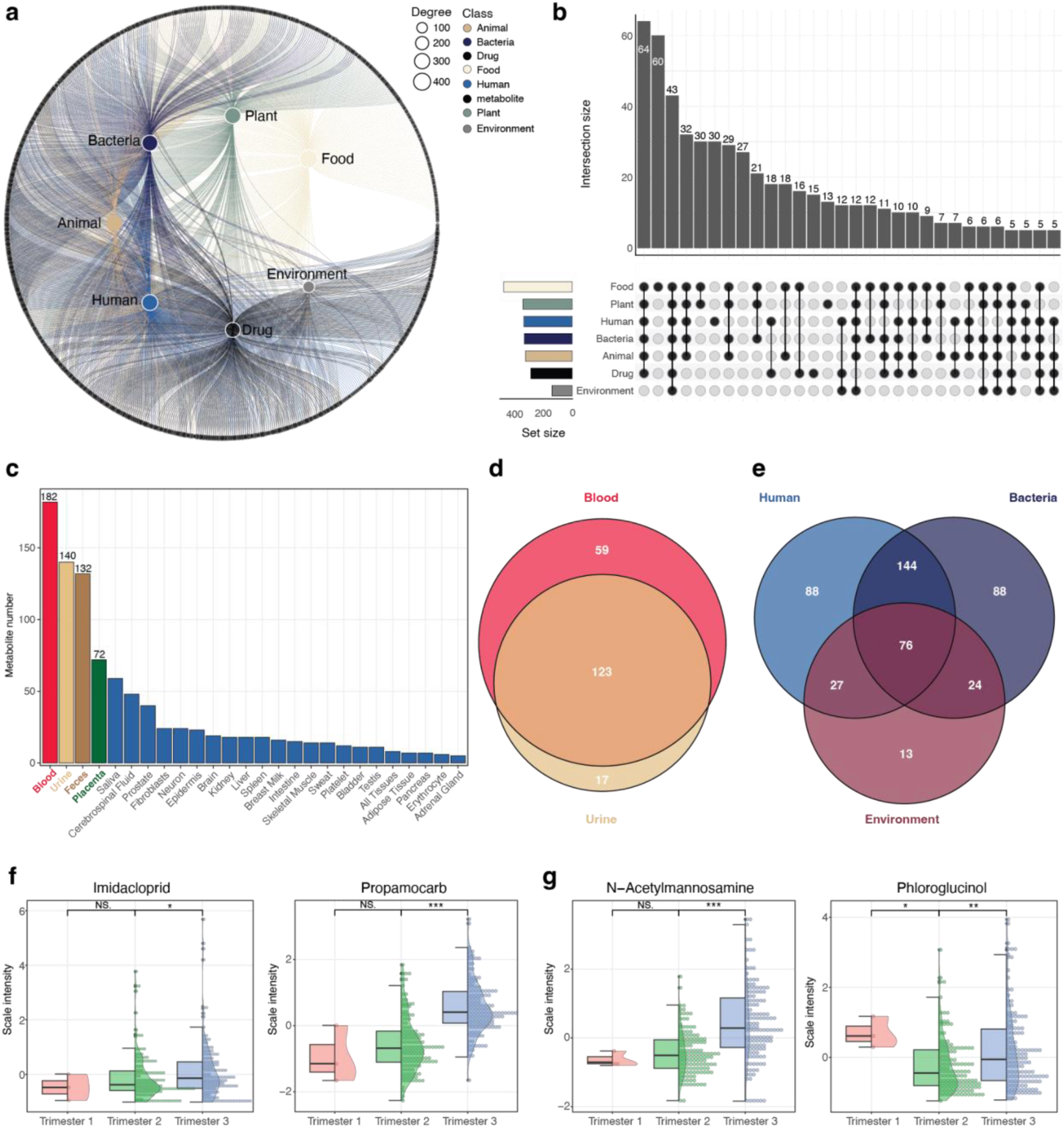
| Diverse origins of urinary metabolites during human pregnancy. **a**, Circular network plot showing the origin-based classification of 662 metabolites. **b,** UpSet plot showing the intersections between major source categories. **c,** Bar plot illustrating the number of pregnancy urine metabolites classified by their specific human-derived sources. Blood, urine, feces, and placenta represent the most frequently assigned origins. **d,** Venn diagram showing the overlap between blood– and urine-derived metabolites, with 123 metabolites (61.8%) shared between the two categories. **e,** Venn diagram highlighting shared and unique metabolites among the human, bacterial, and environmental sources. A substantial number of metabolites were uniquely attributed to each category. **f,** Violin plots showing trimester-specific abundance patterns for two environmental-source metabolites, Imidacloprid and Propamocarb. **g,** Violin plots showing trimester-specific abundance patterns for two microbiome-source metabolites, N-Acetylmannosamine and Phloroglucinol.

We first assessed the distribution of these metabolites across the seven origin categories defined in our database. Among the 662 metabolites, 335 were classified as human-derived (**Figure 4b**). Further resolution of these human-derived metabolites revealed their specific biological sources, with the majority originating from blood, followed by urine, feces, and placenta (**Figure 4c**). This pattern aligns with known physiological processes, as metabolites from these compartments are frequently excreted in urine. A notable overlap was observed between metabolites annotated as blood– and urine-sourced: 123 out of 199 blood-sourced metabolites (61.8%) were also found in urine (**Figure 4d**). This overlap reinforces the value of urine as a surrogate matrix for assessing systemic metabolic alterations during pregnancy.

Next, we focused on three metabolite source categories that are highly relevant to maternal and fetal health: human, environmental, and bacterial. Although substantial overlaps were observed among these categories, a considerable number of metabolites remained unique to each source (**Figure 4e**). This finding underscores the complexity of the maternal metabolic landscape and highlights how urine metabolomics captures a composite of endogenous processes, microbial metabolism, and environmental exposures.

Among the environment-sourced metabolites detected in maternal urine, two compounds, Imidacloprid and Propamocarb, showed significantly elevated levels in the third trimester compared to the second (**Figure 4f**). These two detected environmental contaminants likely entered the maternal system through dietary exposure, as both are commonly used in conventional agriculture^71,72^. Imidacloprid is a neonicotinoid insecticide widely applied to fruits, vegetables, and grains^73,74^, while Propamocarb is a fungicide frequently used on potatoes, lettuce, and tomatoes^75,76^. Some studies have shown that they can be detected in human blood and urine samples^77–79^. Their increasing concentrations during late pregnancy might reflect either accumulation in maternal tissues, changes in maternal detoxification capacity, or seasonal variations in dietary exposure patterns. These findings demonstrate how TidyMass2 can identify potential environmental exposures requiring further investigation, particularly in vulnerable populations like pregnant women.

In addition, TidyMass2 facilitated the detection of microbiome-derived metabolites that increased during late pregnancy. Two such metabolites, N-Acetylmannosamine and Phloroglucinol, were significantly upregulated in the third trimester compared to the second (**Figure 4g**). These metabolites are known microbial products with diverse physiological roles, including modulation of immune tolerance and oxidative balance^80,81^. Their dynamic changes may reflect microbiome-driven metabolic adaptations critical to maintaining pregnancy. Importantly, these metabolites may not have been prioritized using traditional annotation pipelines, highlighting the strength of our origin-informed framework.

In summary, these results highlight the diverse and multi-origin composition of the pregnancy urine metabolome. By integrating comprehensive metabolite origin inference with LC-MS untargeted metabolomics, TidyMass2 enables the discovery of biologically relevant endogenous, microbial, and environmental metabolites. This capacity expands the interpretability of urine metabolomics in pregnancy and offers new avenues for understanding maternal-fetal health, environmental exposures, and host-microbiome interactions during gestation.

### Dysregulated Metabolic Modules During Human Pregnancy

During pregnancy, substantial metabolic remodeling occurs, providing potential biomarkers for pregnancy-related conditions such as preterm birth, gestational diabetes mellitus (GDM), and preeclampsia^82–84^. These metabolic changes also offer insights into the underlying mechanisms of pregnancy and its associated complications^85^. In the longitudinal human pregnancy study, we applied comprehensive urine metabolomics analysis to characterize these changes. Using significance analysis of microarrays (SAM) and correlation analysis, we identified 936 metabolic features that significantly increased and 108 that significantly decreased across pregnancy (**Figure 5a**). To validate the pregnancy-specific nature of these changes, we calculated a recovery score that measures how metabolic features change after delivery compared to late pregnancy (GA range: 28-42 weeks, **Methods**). Remarkably, 98.82% (925 out of 936) of increased metabolic features exhibited recovery scores less than 1, indicating a significant decrease after delivery. Correspondingly, all decreased metabolic features during pregnancy showed recovery scores greater than 1, confirming a return to higher levels postpartum (**Figure 5a** and **Supplementary** Figure 4a). This consistent pattern of reversal strongly suggests that the identified metabolic features represent genuine pregnancy-associated changes, providing a robust foundation for comprehensive metabolic insights.

**Figure 5.**
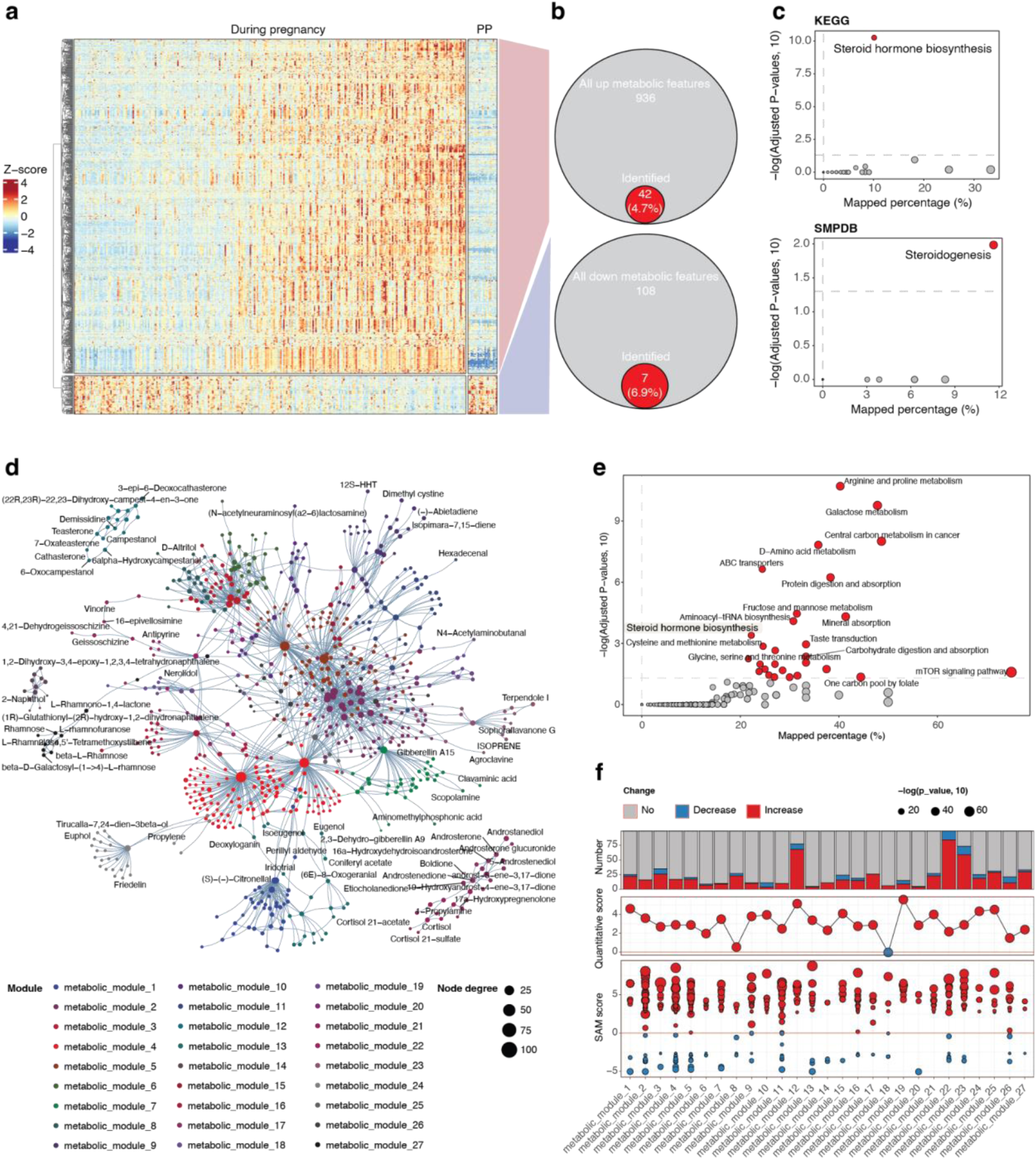
| Comprehensive metabolic module analysis reveals extensive metabolic reprogramming during pregnancy. **a**, Heatmap showing the dynamic changes of significant metabolic features across pregnancy and postpartum (PP). Z-scores indicate relative abundance changes, with red representing increased and blue representing decreased levels. **b,** Pie charts illustrating the annotation rate of significantly changing metabolic features during pregnancy. **c,** Pathway enrichment analysis using MS^2^ spectra-based annotated metabolites. **d,** Network visualization of the 27 dysregulated metabolic modules identified using TidyMass2’s feature-based module and pathway analysis. Nodes represent metabolites (labeled), with node size indicating degree of connectivity. Colors indicate distinct metabolic modules. **e,** Enhanced pathway enrichment analysis using all metabolites in the dysregulated metabolic network, showing 29 enriched pathways with higher mapped percentages compared to traditional approaches. **f,** Quantitative characterization of the 27 metabolic modules.

Despite the large number of significant metabolic features identified, traditional MS^2^ spectra-based metabolite annotation approaches yielded limited results; only 4.7% (42 out of 936) of increased metabolic features and 6.9% (7 out of 108) of decreased metabolic features were successfully annotated (**Figure 5b**). Pathway enrichment analysis using these annotated metabolites and referencing KEGG and SMPDB databases revealed steroid hormone biosynthesis (KEGG) and steroidogenesis (SMPDB) as significantly enriched pathways, consistent with previous studies^86,87^. These two pathways refer to the same pathway as defined in both databases (**Supplementary** Figure 4b). However, the low annotation rate severely restricted our ability to comprehensively understand metabolic changes during pregnancy.

To decode the comprehensive metabolic change during pregnancy, we applied the metabolic feature-based functional module analysis approach in TidyMass2, which significantly expands biological interpretation beyond MS^2^ spectra-based annotated metabolites. Finally, this approach identified 27 dysregulated metabolic modules (**Supplementary Table 3**), creating a dysregulated metabolic network that incorporated a substantially higher percentage of the significantly changing features compared to traditional methods (**Figure 5d)**. This network-based approach enabled the utilization of more than 58.8% (614 out of 1,044) of significantly changed features for biological insight mining (**Supplementary** Figure 4c), representing a marked improvement over traditional annotation-dependent methods, which only utilized 5.8% of significant metabolic features (**Figure 5b**).

Then we performed pathway enrichment analysis using all metabolites in the dysregulated metabolic network; 29 pathways were enriched (**Figure 5e, Supplementary Table 4**), a significant expansion compared to the limited pathways identified through traditional methods (29 *vs.* 1 pathways) (**Figure 5c**). The enriched pathways encompassed broader aspects of metabolism, including lipid metabolism, amino acid processing, and energy production pathways relevant to pregnancy.

We then carried out quantitative analysis of the dysregulated modules throughout pregnancy, which revealed that most metabolic modules increased during pregnancy (**Figure 5f**), consistent with the pattern observed in individual metabolic features (**Figure 5a**). As expected, most of these metabolic modules showed increasing temporal patterns, with some exhibiting early changes in the first trimester while others progressively increased across gestation (**Supplementary** Figure 5a). Module correlation analysis further revealed interconnected positive correlation relationships between them (**Supplementary** Figure 5b), suggesting coordinated metabolic reprogramming to support fetal development and maternal adaptation.

To gain deeper biological insights from our metabolic network-based analysis, we further characterized several key dysregulated metabolic modules with distinct functional significance (**Figure 6**). The feature-based module analysis enabled us to link unidentified metabolic features with metabolic pathways (**Methods**), substantially expanding our ability to extract biological insights from the dataset.

**Figure 6.**
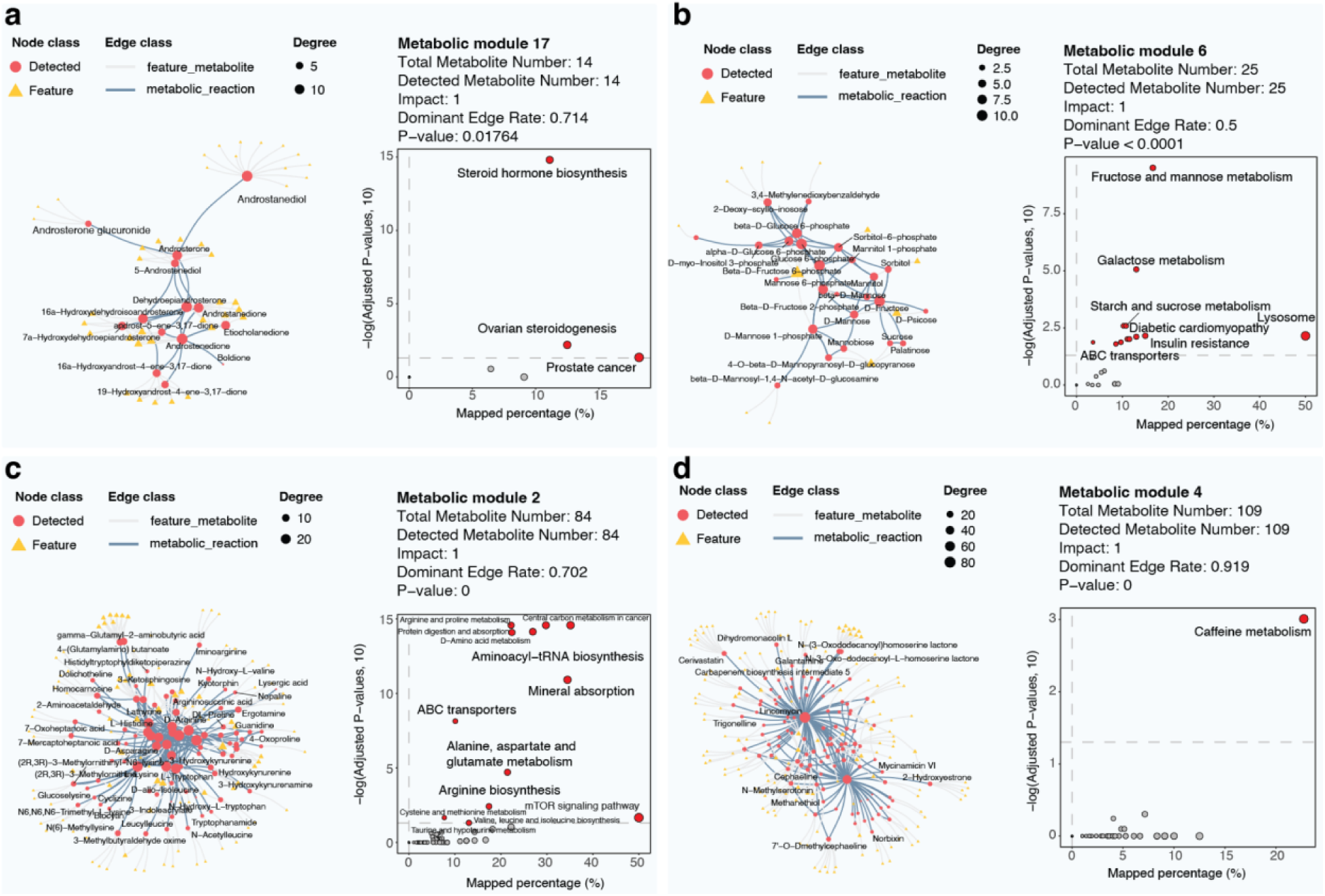
| Functional characterization of key dysregulated metabolic modules identified in maternal urine during pregnancy. **a**, Metabolic module 17 is associated with steroid hormone pathways. **b,** Metabolic module 6 representing carbohydrate metabolism. **c,** Metabolic module 2 is associated with amino acid metabolism. **d,** Metabolic module 4 representing caffeine metabolism. Left: Network visualization showing metabolic features (yellow triangles) and detected metabolites (red circles). Node size indicates the degree of connectivity. Right: Pathway enrichment analysis of the metabolites on the dysregulated metabolic modules.

The metabolic module 17 comprised 57 metabolic features and 14 metabolites, which is primarily associated with steroid hormone-related pathways, including steroid hormone biosynthesis (p-value < 0.001) and ovarian steroidogenesis (p-value < 0.001) (**Figure 6a**). The module contained multiple androgen derivatives and glucuronides, highlighting the capacity of our approach to capture established pregnancy-associated endocrine changes^88,89^. The enrichment of steroid-related pathways is consistent with previous studies on maternal metabolic adaptations during pregnancy^90,91^.

Metabolic module 6 consisted of 16 metabolic features and 25 metabolites focused primarily on carbohydrate metabolism (**Figure 6b**). This module revealed significant enrichment in fructose and mannose metabolism (p-value < 0.001), galactose metabolism (p-value < 0.001), and starch and sucrose metabolism pathways (p-value < 0.001). Interestingly, this module also showed connections to disease-relevant pathways, including diabetic cardiomyopathy (p-value < 0.001) and insulin resistance (p-value < 0.001), potentially reflecting the metabolic stress associated with pregnancy and the increased risk for gestational diabetes mellitus^92,93^.

One of the most extensive dysregulated networks, metabolic module 2, contained 133 metabolic features and 84 metabolites (**Figure 6c**). This module was predominantly associated with amino acid metabolism, showing significant enrichment in aminoacyl-tRNA biosynthesis (p-value < 0.001), alanine, aspartate and glutamate metabolism (p-value < 0.001), arginine biosynthesis (p-value < 0.001), and cysteine and methionine metabolism (p-value < 0.05). Additionally, this module is linked to important cellular processes, including ABC transporters (p-value < 0.001), mineral absorption (p-value < 0.001), and mTOR signaling (p-value < 0.005), which reflect the increased protein synthesis and nutrient transport demands during pregnancy.

In metabolic module 4, we identified a dysregulated metabolic network containing 208 metabolic features and 208 metabolites (**Figure 6d**). This module was significantly associated with caffeine metabolism (p-value < 0.001), a finding that aligns with and confirms observations from parallel plasma sample analyses, providing cross-validation between different biofluids^91^.

Together, these results demonstrate that the metabolic feature-based functional module analysis approach implemented in TidyMass2 successfully identified both well-established pregnancy-associated metabolic changes and novel biological insights. The approach validated findings across different biological matrices (urine and plasma) while uncovering previously uncharacterized metabolic adaptations during pregnancy. By integrating unannotated metabolic features into biologically meaningful modules, our method substantially expanded the interpretative scope beyond traditional MS^2^ spectra-based metabolite annotation analyses, enabling more comprehensive metabolic phenotyping during pregnancy.

## Discussion

In this study, we present TidyMass2, a comprehensive computational framework that addresses critical challenges in LC-MS untargeted metabolomics data analysis. The three major innovations implemented in TidyMass2: metabolite origin inference, metabolic feature-based functional module analysis, and an intuitive web interface, significantly advance our ability to derive meaningful biological insights from complex untargeted metabolomic datasets, particularly in contexts where conventional annotation-dependent approaches provide limited information.

The metabolite origin inference system represents a major step forward in understanding the complex interplay between endogenous metabolism, the microbiome, diet, pharmaceuticals, and environmental exposures. By integrating source information from 11 distinct databases, we created the MetOriginDB and a unified reference system that enables researchers to trace the origins of detected metabolites. The substantial overlap we observed between human and bacterial metabolites provides quantitative confirmation of the extensive metabolic integration between host and microbiome. This finding underscores the concept of “co-metabolism,” where metabolic products arise from multi-organism metabolic pathways. The ability to distinguish between endogenous and exogenous metabolites provides a critical context for interpreting results, particularly in studies investigating host-microbiome interactions, exposome research, and nutritional interventions.

Another important advancement in TidyMass2 is the metabolic feature-based functional module analysis approach. Traditional metabolomics studies face a fundamental limitation in that typically, only ∼10%-20% of detected metabolic features can be annotated using MS^2^ spectral matching, leaving the vast majority of potentially valuable biological information inaccessible^14^. Our approach leverages the inherent network structure of metabolism to extract biological meaning from unannotated features, dramatically expanding the interpretative scope of metabolomics data. The pregnancy case study demonstrates the power of this approach, where we increased the proportion of biologically interpretable features from 5.8% to 58.8% and expanded the number of enriched pathways from just 1 to 29. This substantial improvement enables more comprehensive metabolic phenotyping and mechanistic understanding.

The comprehensive human metabolic network we constructed, comprising 9,630 metabolites connected by 30,196 edges, represents the largest human metabolic network currently available to our knowledge^94,95^. This resource not only underpins the metabolic feature-based functional module analysis in TidyMass2 but also serves as a valuable reference for the broader metabolomics community. The network’s scale-free properties, with most metabolites having connectivity degrees between 1 and 10, align with established characteristics of biological networks and provide confidence in its biological relevance^96,97^.

The entirely open-source nature of TidyMass2, with all code and databases freely available, promotes reproducibility and transparency in untargeted metabolomic data analysis. The addition of an intuitive Shiny interface significantly lowers the technical barriers to utilizing advanced metabolomics analysis tools, making sophisticated computational approaches accessible to researchers without extensive programming expertise. This commitment to accessibility aligns with broader movements toward open science and democratization of advanced analytical methods.

Compared to existing metabolomics analysis tools^17,23,24,37,38^, TidyMass2 offers several distinctive advantages (**Supplementary Table 2**). While these platforms provide robust solutions for specific aspects of metabolomics analysis, TidyMass2 uniquely integrates metabolite origin inference with feature-based functional module analysis in a comprehensive framework. For example, MetaboAnalyst offers statistical and pathway analysis but lacks metabolite origin inference capabilities and customized databases and pathway databases for metabolite annotation and pathway enrichment. MS-DIAL focuses on raw data processing and metabolite annotation but does not leverage the metabolic feature-based functional module. TidyMass2 addresses these limitations by offering an end-to-end and comprehensive solution that maintains data integrity throughout the analysis workflow while significantly expanding biological interpretation.

The application of TidyMass2 to urine metabolomics data from human pregnancy revealed extensive metabolic remodeling with high pregnancy specificity, as evidenced by the consistent reversal patterns observed in recovery scores. The identification of 27 dysregulated metabolic modules provides a systems-level view of pregnancy-associated metabolic changes, capturing well-established pathways such as steroid hormone biosynthesis while uncovering previously uncharacterized adaptations in carbohydrate and amino acid metabolism. The cross-validation of findings between urine and plasma samples, as demonstrated with caffeine metabolism, further supports the robustness of our approach^98^. The identification of caffeine metabolism as a significantly dysregulated pathway during pregnancy is particularly interesting given known alterations in caffeine pharmacokinetics during gestation^91^. Pregnancy extends the half-life of caffeine from approximately 3 hours in non-pregnant women to 10-16 hours by late pregnancy, primarily due to decreased CYP1A2 activity^99^. These metabolic changes may explain our observations and highlight the importance of caffeine monitoring during pregnancy, given its potential associations with fetal growth and development outcomes^100^.

Beyond its application to pregnancy metabolomics demonstrated in this study, TidyMass2 is designed to benefit diverse research domains, including host-microbiome interactions, environmental exposomics, pharmacometabolomics, and nutritional metabolomics. The ability to trace metabolite origins and extract biological meaning from unannotated features makes TidyMass2 particularly valuable for complex, multi-factorial studies where conventional annotation-dependent approaches provide limited insights.

Despite these advances, several limitations and opportunities for future development remain. First, while our metabolite origin database represents a substantial improvement over existing resources, the classification system remains relatively broad. Future refinements could incorporate more granular origin categories, particularly for microbiota-derived metabolites, where species-level attribution could provide valuable insights into specific microbial contributions. Second, the metabolic feature-based functional module analysis, while powerful, still relies on putative mass-based annotations as a starting point. Integration with emerging computational MS^2^ spectra prediction methods could further improve annotation confidence and expand coverage. Third, the current implementation focuses primarily on human metabolism, and extension to other model organisms would broaden applicability across diverse research domains. Addressing these limitations represents our roadmap for future development of the TidyMass ecosystem.

In conclusion, TidyMass2 represents a significant advancement in computational metabolomics, addressing key limitations in metabolite origin determination and biological interpretation based on limited MS^2^ spectra-based annotated metabolic features. The framework’s ability to extract meaningful biological insights from incompletely annotated datasets substantially expands the utility of untargeted metabolomics across diverse biomedical applications. As multi-omics approaches continue to transform biomedical research, TidyMass2 provides a powerful tool for integrating metabolomic data into comprehensive biological understanding, ultimately contributing to improved mechanistic insights, biomarker discovery, and therapeutic development.

As a long-term project, our roadmap for TidyMass development focuses on addressing current limitations and expanding capabilities. Near-term objectives include: (1) implementing more granular origin classification with species-level resolution for microbiome-derived metabolites; (2) integrating emerging computational MS^2^ spectra prediction methods to improve annotation confidence; (3) expanding support for non-human organisms by incorporating additional model organism metabolic networks; and (4) developing enhanced capabilities for multi-omics data integration, such as microbiome and metabolome, and exposome and metabolome. Additionally, community contributions are encouraged through our GitHub repository, where feature requests and bug reports can be submitted. Regular updates will be provided through the TidyMass website (https://www.tidymass.org).

## Methods

### TidyMassShiny development

*TidyMassShiny* was developed as an integral component of TidyMass2 to provide a user-friendly graphical interface, making advanced metabolomics data analysis accessible to researchers regardless of their computational background. The application was built using R’s Shiny framework, which enables interactive web applications directly from R code, allowing us to create a seamless bridge between powerful analytical capabilities and an intuitive user experience.

The design philosophy of *TidyMassShiny* centers on modularity, reproducibility, and interoperability. We implemented a comprehensive data processing pipeline that guides users through the entire metabolomics workflow, from raw data processing and quality control to advanced analyses, including metabolite annotation, origin tracing, and pathway enrichment. Each module corresponds to a key step in the TidyMass2 framework, preserving the analytical rigor while abstracting away programming complexity.

A distinctive feature of *TidyMassShiny* is its seamless integration with the “mass_dataset” object structure fundamental to TidyMass2. Users can upload mass_dataset objects at any stage of the workflow, allowing entry at any point in the analysis pipeline. This flexibility enables researchers to perform certain steps using alternative tools and then continue their analysis within *TidyMassShiny* without data format conflicts. Similarly, at each analytical stage, the “mass_dataset” object is automatically saved. This feature facilitates the reproducibility of analysis steps, integration with external tools, and support for collaborative workflows across research teams utilizing diverse software preferences.

The interface design prioritizes scientific visualization, providing interactive plots for data exploration, quality assessment, and result interpretation. We implemented customizable visualization options for multiple data types, including peak intensity distributions, principal component analyses, metabolic networks, and pathway enrichment results. These visualizations are generated dynamically, allowing researchers to adjust parameters and immediately observe the effects on their data.

While *TidyMassShiny* offers substantial analytical power, its extensive dependency framework, requiring numerous R packages for specialized statistical, chemometric, and visualization functions, created significant installation challenges, particularly for researchers without computational expertise. To address this barrier, we developed a containerized Docker version of *TidyMassShiny* that encapsulates the entire application with all dependencies in a pre-configured environment.

The Docker implementation provides several advantages: (1) it eliminates dependency conflicts and version compatibility issues that frequently arise during manual installation; (2) it ensures consistent performance across operating systems, including Windows, macOS, and Linux distributions; and (3) it requires minimal setup knowledge, as users need only install Docker and execute a simple command to retrieve and run the *TidyMassShiny* image.

We optimized the Docker image for both performance and accessibility, carefully balancing resource utilization with processing speed. The container includes pre-installed versions of all required R packages, optimized computational libraries, and a complete runtime environment configured specifically for metabolomics analysis. This approach maintains analytical performance while significantly reducing the technical barriers to adoption.

For server deployments, we included configuration options that enable institutions to deploy *TidyMassShiny* on their internal networks, supporting multiple concurrent users while maintaining data security and privacy. The Docker image is regularly updated to incorporate bug fixes, feature enhancements, and security updates, with versioning that ensures reproducibility of analyses across time. This containerized approach has substantially expanded TidyMass2’s accessibility, enabling researchers from diverse backgrounds, including those with limited programming experience, to leverage advanced metabolomics analysis methods previously restricted to those with specialized computational training.

Computational performance was a key priority in the development of *TidyMassShiny*. The application was optimized to efficiently process large-scale metabolomics datasets while maintaining responsive performance on standard hardware. To address the challenges of scalability, *TidyMassShiny* is completely open source and supports local server deployment as we described above, enabling users to leverage their own computational resources. This approach offers a sustainable solution, as providing centralized cloud computing power for all users would be impractical and unsustainable in the long term.

To ensure reliability and reproducibility, TidyMass2 was rigorously tested across multiple platforms and datasets. Core functions were validated using unit tests implemented with the *testthat* framework. All TidyMass2 packages are actively maintained by our team and the broader user and developer community via GitHub, enabling continuous updates, issue tracking, and collaborative improvements.

### Metabolite database with source information for metabolite origin inference

TidyMass2 integrates 11 diverse metabolite databases with source information to create a comprehensive metabolite reference system. These databases include KEGG^29^, HMDB^28^, BiGG models^43^, DrugBank^31^, ChEBI^42^, FooDB (www.foodb.ca), LOTUS^46^, MiMeDB^44^, ModelSEED^47^, Reactome^36^, and T3DB^45^ (**Figure 2a**). These 11 databases were selected based on their complementary coverage of different metabolite sources, established reputation in the field, and availability of structured data for integration. Each database contributes unique strengths, and for instance, HMDB provides comprehensive human metabolite coverage with detailed biofluid information; KEGG offers pathway context across diverse organisms; MiMeDB specializes in microbial metabolites with species-level resolution; and T3DB focuses specifically on environmental toxins and contaminants.

We systematically categorized all metabolites into seven source classes: human, animal, bacteria, drug, food, plant, and environment (**Figure 2a**). Each metabolite in the database is systematically classified using a standardized annotation schema. If a metabolite is of human origin, its “from_human” column is designated as “Yes”; if evidence indicates it is not from human sources, this column is marked as “No”; and in cases where origin information is unavailable or inconclusive, the value is recorded as “Unknown.” This consistent three-tier classification system (Yes/No/Unknown) is applied across all seven source categories, enabling precise filtering and comprehensive metabolite origin inference. For each source category, we implemented detailed specificity annotations where available. For instance, human-derived metabolites are further characterized by their biological origin. For example, if a metabolite is found in human blood, it receives a “blood” designation in the “from_which_part” column. Similarly, bacterial metabolites are annotated with their specific microbial source, such as a metabolite produced by Escherichia coli being marked as “e.coli” in the “from_which_bacteria” column. This granular classification system preserves valuable provenance information, enabling a more nuanced interpretation of metabolite origins in complex biological samples.

*KEGG.* The Kyoto Encyclopedia of Genes and Genomes (KEGG) contains metabolic pathway information spanning thousands of species^101^. Using the *massdatabase* package, we downloaded and analyzed all available metabolic pathways, extracting and classifying metabolites according to their taxonomic origins. Metabolites present in human pathways were labeled as human-derived, while those in bacterial pathways received bacterial classification. Additionally, we incorporated the KEGG DRUG database, labeling all entries as drug-sourced compounds. The final KEGG integration yielded approximately 28,804 metabolites and 11,873 drug compounds with source annotations.

*HMDB.* The Human Metabolome Database (HMDB) provides extensive information on metabolites detected in various human biological samples^28^. We processed all 217,920 HMDB metabolites, assigning human source classification by default. Where origin filter options such as “Food”, “Plant”, and “Microbial” were provided in the HMDB website, we adopted the labels for the corresponding metabolites and replaced the original human source. HMDB’s biofluid and cellular localization data provided additional context regarding metabolite distribution within human systems.

*BiGG models.* The Biochemical Genetic and Genomic knowledgebase (BiGG) contains metabolic models across diverse species. We systematically downloaded information for each organism, extracted associated metabolites, and assigned source classifications based on taxonomic information. This approach enabled multi-source labeling; metabolites present in both bacterial and plant models, for example, received both source designations. BiGG’s curated genome-scale metabolic reconstructions provided reliable species associations for metabolites. The final database contains 7,027 metabolites.

*DrugBank.* This comprehensive pharmaceutical database contains detailed molecular and pharmacological data on thousands of drug compounds^31^. We integrated the complete DrugBank dataset, classifying all metabolites as drug-sourced. This integration enhances TidyMass2’s ability to distinguish pharmaceutical compounds from endogenous metabolites in biological samples. The final database contains 11,300 metabolites.

*ChEBI.* The Chemical Entities of Biological Interest (ChEBI) database provides a structured classification of molecular entities with biological relevance. We leveraged ChEBI’s “Metabolite of Species” ontology to extract taxonomic origin information for each metabolite, enabling precise source classification across kingdoms of life. ChEBI’s ontological structure facilitated the hierarchical classification of metabolite origins. The final database contains 147,390 metabolites.

*FooDB.* The Food Database (FooDB) catalogs the chemical composition of common unprocessed foods. For each metabolite, FooDB specifies food sources, allowing us to label all entries as food-derived while preserving specific information about particular food origins (*e.g.*, specific fruits, vegetables, or animal products). This fine-grained food source information provides valuable context for nutritional metabolomics studies. The final database contains 70,360 metabolites.

*LOTUS.* The naturaL prOducTs occUrrence databaSe (LOTUS) specializes in natural products with detailed source information^46^. We classified metabolites according to their documented origins, primarily spanning plant, fungal, and bacterial sources, enhancing coverage of naturally occurring bioactive metabolites. LOTUS’s taxonomically organized data structure enabled the precise attribution of metabolites to specific biological sources. The final database contains 276,518 metabolites.

*MiMeDB.* The Microbial Metabolites Database (MiMeDB) focuses specifically on small molecules produced by the human microbiome. We incorporated all metabolites from this database with bacterial source classifications, significantly enriching TidyMass2’s coverage of gut microbiome-derived metabolites. MiMeDB’s focus on human microbiome metabolites provides critical information for studies investigating host-microbiome metabolic interactions. The final database contains 27,641 metabolites.

*ModelSEED.* ModelSEED is a biochemistry database containing metabolites associated with 41 plant species. We downloaded all plant-related data, extracted the metabolites, and labeled them according to their plant origins, including specific plant species information where available. This addition significantly enhanced our coverage of plant metabolites with precise taxonomic sourcing, which is particularly valuable for studies involving dietary plant metabolites. The final database contains 33,992 metabolites and 512 metabolites with plant source information.

*Reactome.* Reactome provides metabolic pathways for different species, including humans and various microorganisms. We downloaded the mapping files of ChEBI to all pathways, extracted metabolites, and integrated them into our metabolite database with corresponding source information. Reactome’s pathway context provided additional functional context for metabolites, complementing the source attribution data. The final database contains 2,487 metabolites.

*T3DB.* The Toxin and Toxin Target Database (T3DB) is a specialized bioinformatics resource that combines detailed toxin data with comprehensive toxin target information. We downloaded all 3,678 metabolites and classified them as environment-sourced metabolites. T3DB’s focus on environmental toxins enhances TidyMass2’s ability to identify potentially harmful exogenous metabolites in metabolomic profiles.

*Integration of the comprehensive metabolite database.* All 11 metabolite databases were integrated into a unified and comprehensive metabolite database with source information. During integration, we employed a hierarchical identifier matching strategy, prioritizing HMDB identifiers first, followed by KEGG identifiers, InChI strings, InChIKeys, and CAS registry numbers. For source information reconciliation, we implemented a knowledge-preserving approach: if a metabolite was labeled with a specific source in one database but had an uncertain origin in another database. We retained the specific source classification. We employed a stepwise pairwise integration approach, first combining two databases and then incorporating additional databases sequentially to ensure maximal coverage and accuracy. The final database (MetOriginDB) was implemented as a database class in the *metpath* package. It is entirely open-source and available from the TidyMass website: https://www.tidymass.org/databases/. Finally, there were 827,117 metabolites in total for the 11 databases, and the comprehensive metabolite database contains over 650,884 unique metabolites, and 532,488 of them have at least one source annotation.

*MS*^2^ *spectra databases with source information.* We enhanced the utility of our MetOriginDB by integrating source information with our MS^2^ spectral databases, encompassing both in-house and public MS^2^ spectra databases^21,28,102^. This integration enables source attribution during metabolite annotation via spectral matching, a key innovation that distinguishes TidyMass2 from existing tools. This MS^2^ spectra-enhanced database significantly improves the specificity and accuracy of metabolite source attribution in untargeted metabolomics studies. Finally, 25 MS^2^ spectra databases were created with metabolite source information **(Supplementary** Figure 3a,b), which contains 81,461 metabolites and 502,790 MS^2^ spectra (MetOriginDB_ms2; 342,892 for positive mode and 159,898 for negative mode). And all the MS^2^ spectra databases are open source and can be downloaded from the TidyMass website: https://www.tidymass.org/databases/.

### Metabolite origin inference

TidyMass2 offers two complementary approaches for metabolite origin inference. In the first approach, users can employ the integrated *metid* package for metabolite annotation^27^, and then seamlessly analyze origin information by passing their annotated “mass_dataset” object to the “analyze_metabolite_origin” function. Alternatively, for increased flexibility, users who prefer external annotation tools can import their independent annotation results into the “analyze_metabolite_origin” function. This dual-pathway design accommodates diverse workflows while providing consistent access to TidyMass2’s comprehensive metabolite origin database, enabling researchers to trace the biological, dietary, pharmaceutical, or environmental sources of detected metabolites regardless of their preferred annotation methodology.

### Metabolic network

The metabolic network in TidyMass2 was constructed by integrating reaction pair information from eight public databases: ModelSeed^47^, Recon3^103^, KEGG^29^, MetaNetx^104^, BiGG^43^, Rhea^105^, HMDB^28^, and Reactome^36^. All reaction pair data were retrieved using the *massdatabase* package^26^ and subsequently merged, with duplicate reaction pairs removed across databases (**Figure 2d**). The resulting consolidated network comprises 9,630 nodes (metabolites) and 30,196 edges (reactions). This network was structured as a network file (tbl_graph class) using the *igraph* and *ggraph* packages in R. The complete metabolic network is available for download at https://www.tidymass.org/databases/, while the code for downloading reaction pairs and data processing is accessible at https://github.com/tidymass/metabolomics_databases.

### Metabolic feature-based functional module analysis

The feature-based functional module analysis implemented in TidyMass2 expands traditional pathway analysis beyond annotated metabolites, addressing the annotation bottleneck in untargeted metabolomics. This approach, available in the *metpath* package and *TidyMassShiny* interface, provides comprehensive insights into metabolic changes. Detailed documentation is available at https://www.tidymass.org/tidymassshiny-tutorial/.

*Data preparation.* Users must provide two feature tables: (1) a comprehensive table containing all metabolic features with variable_id (unique identifier), *m/z* (mass-to-charge ratio), rt (retention time in seconds), and polarity information (“positive” or “negative”); and (2) a marker feature table of statistically significant features that additionally includes log2(fold change) or correlation values and p-values. The analysis also requires a metabolic network, metabolite database, and pathway database, all available for download from the TidyMass2 website.

*Metabolite annotation based on m/z.* Initially, all metabolic features are matched to potential metabolites in the database using MS1-based annotation through the *metid* package. With a default mass tolerance of 25 ppm (user-adjustable), all annotations within the threshold are retained for subsequent analysis.

*Isotope annotation.* For each metabolite-feature match, we calculate theoretical isotope patterns and search for corresponding features in the feature table. These isotope annotations are incorporated into the annotation table, providing additional confidence for metabolite identifications.

*Metabolite class identification.* Annotations are organized into metabolite classes based on chemical identity and chromatographic behavior. A metabolite class comprises metabolic features with identical annotations and similar retention times (within a defined tolerance). This approach recognizes that one metabolite may form multiple metabolite classes due to retention time differences, while a single feature may belong to several metabolite classes with different annotations.

*Scoring for metabolite class.* Each metabolite class receives a confidence score based on adduct and isotope pattern evidence. The scoring system awards points for characteristic adducts in each ionization mode: 50 points for primary adducts ([M+H]+ in positive mode or [M-H]-in negative mode), 20 points for isotopes of these primary adducts, 20 points for alternative adducts, and 10 points for isotopes of alternative adducts. The maximum possible score is 200, with higher scores indicating greater annotation confidence.

*Removing redundant annotations based on scores.* To reduce false positives, we filter annotations based on confidence scores. When a feature belongs to multiple metabolite classes, if the highest-scoring class exceeds 100 points, lower-scoring alternatives are removed. This process significantly reduces annotation redundancy and eliminates likely incorrect assignments.

*Identifying hidden metabolites from the metabolic network.* To enhance pathway coverage, we incorporate “hidden metabolites” that connect detected metabolites within the reaction network. Hidden metabolites are defined as metabolites that are not directly detected but are capable of connecting any two detected metabolites within three reaction steps. These hidden metabolites increase network integrity, which is particularly valuable when working with a limited number of detected metabolites.

*Identifying dysregulated metabolic networks and modules.* Using both detected and hidden metabolites, we extract a subnetwork from the comprehensive metabolic network, termed the dysregulated metabolic network. The community detection algorithms implemented in TidyMass2 leverage random walks through the metabolic network to identify densely connected subgraphs that likely represent functional modules. Specifically, we employ the Walktrap algorithm, which uses the principle that short random walks tend to stay within the same community due to higher internal edge density^106^. This approach has been validated in protein-protein interaction networks and metabolic networks, showing superior performance in identifying biologically meaningful modules compared to alternative methods^107,108^.

Compared to existing network-based approaches like Mummichog^23^ and PIUMet^24^, our metabolic feature-based functional module analysis offers several advantages. Unlike Mummichog, which cannot integrate data from positive and negative ionization modes, TidyMass2 processes data from both modes simultaneously, increasing metabolite coverage. Our approach also incorporates retention time information, which substantially reduces annotation redundancy compared to PIUMet’s m/z-only approach. Additionally, by employing a comprehensive scoring system for metabolite class identification, TidyMass2 provides more granular confidence assessment for putative annotations.

*Functional annotation for dysregulated metabolic modules.* Significantly dysregulated modules (p-value < 0.05) were subjected to pathway enrichment analysis using an integrated pathway database containing KEGG, WikiPathways, and Reactome pathways. This analysis revealed biological functions associated with each metabolic module, connecting network-level observations to established biological pathways.

*Data visualization of results.* The dysregulated metabolic modules and networks are visualized through customized network plots that highlight connections between detected metabolites and hidden metabolites and their organization into functional modules. Additional visualizations of enriched pathways provide biological context for interpreting the network structures in relation to pregnancy progression.

### Sample preparation and analytical conditions for the case study

This study reanalyzed data from a previously published investigation^70^, with no new LC-MS metabolomics data generated. Comprehensive details of sample preparation and analytical conditions are available in our previous publication^70^.

In brief, we analyzed 346 urine samples collected longitudinally from 36 ethnically diverse women throughout pregnancy (11.8-40.7 weeks) and postpartum. Gestational age was determined following the guidelines of the American Congress of Obstetricians and Gynecologists. All samples were random spot urine collections rather than 24-hour collections.

For sample preparation, urine samples were thawed and centrifuged at 17,000 rcf for 10 minutes. We diluted 250 μL of supernatant with 750 μL internal standard mixture, vortexed for 10 seconds, and centrifuged at 17,000 rpm for 10 minutes at 4 °C. The resulting supernatant was used for LC-MS analysis.

A pooled quality control (QC) sample was injected every 10 samples to monitor retention time stability and signal intensity.

Data processing began with the conversion of MS raw data (.raw) to .mzXML (MS1) and .mgf format (QC MS^2^ spectra data) using ProteoWizard software. The .mzXML files were organized into three categories (“Blank,” “QC,” and “Subject”) before peak detection and alignment using the *metprocesser* package. Peak detection and alignment utilized the centWave algorithm with the following parameters: method = “centWave”; ppm = 15; snthr = 10; peakwidth = c(5, 30); snthresh = 10; prefilter = c(3, 500); minifrac = 0.5; mzdiff = 0.01; binSize = 0.025; and bw = 5. This generated an MS1 peak table containing m/z, retention time, and peak abundances.

Data cleaning was performed using a *masscleaner*. We removed peaks detected in fewer than 20% of QC samples and excluded samples with more than 50% missing values. The remaining missing values were imputed using the k-nearest neighbors algorithm. Peak intensities were normalized by dividing by the mean peak intensity to minimize intra-batch analytical variations. Finally, we performed batch correction using the ratio of mean peak values between batches as correction factors.

Metabolite annotation was performed using the *metid* package^27^. Annotations made using our in-house database achieved level 1 confidence according to Metabolomics Standards Initiative (MSI) criteria^109^, while those made using public databases were classified as level 2^27^.

## Data Availability

The LC-MS metabolomics data analyzed in this study were obtained from a previously published study^70^.

## Code Availability

This study utilized R version 4.4.1 and its associated packages for software development, data processing, and statistical analysis. The complete source code for TidyMass2 is freely available at https://github.com/tidymass. All the databases (metabolites, pathways, and metabolic network) are freely available at https://www.tidymass.org/databases/. All code used for data processing, analysis, and visualization in this study can be accessed at https://github.com/tidymass/tidymass2_manuscript.

## Acknowledgments

This work was supported by start-up funding provided to Dr. Xiaotao Shen from the Lee Kong Chian School of Medicine (LKCMedicine) and the School of Chemistry, Chemical Engineering, and Biotechnology (CCEB) at Nanyang Technological University, Singapore (NTU).

## Author Contributions

X.S. conceived the method and provided overall supervision. X.S., Y.L., and X.W. jointly developed the methods, databases, and software packages. X.W. and X.S. implemented *TidyMassShiny* and created its Docker version. Y.L., X.S., and X.W. prepared documentation and tutorials. Y.L. and X.S. analyzed the case study data. X.S. and Y.L. generated all figures. X.S., Y.L., and P.G. wrote the manuscript. All authors reviewed and contributed to the final manuscript.

## Competing Interests

The authors declare no conflict of interest.

## Additional information

Correspondence and requests for materials should be addressed to X.S.

## Notes

https://github.com/tidymass

### Competing Interest Statement

The authors have declared no competing interest.

