## Supplementary materials for "TidyMass2: Advancing LC-MS Untargeted Metabolomics Through Metabolite Origin Inference and Metabolic Feature-based Functional Module Analysis"

**Supplementary Table 1. Databases used in TidyMass2**

| Database | Link | Metabolite source information | $MS^2$ spectra | Pathway | Chemical reaction |
|---|---|---|---|---|---|
| KEGG | https://www.genome.jp/kegg/ | Yes | No | Yes | Yes |
| HMDB | https://www.hmdb.ca/ | Yes | Yes | Yes | Yes |
| Reactome | https://reactome.org/ | Yes | No | Yes | Yes |
| BIGG | http://bigg.ucsd.edu/ | Yes | No | No | Yes |
| CHEBI | https://www.ebi.ac.uk/chebi/ | Yes | No | No | No |
| DrugBank | https://go.drugbank.com/ | Yes | No | No | No |
| FooDB | https://foodb.ca/ | Yes | No | No | No |
| Lotus | https://lotus.naturalproducts.net/ | Yes | No | No | No |
| MIMEDB | https://mimedb.org/ | Yes | No | No | No |
| Modelseed | https://modelseed.org/ | Yes | No | No | Yes |
| T3DB | https://www.t3db.ca/ | Yes | No | No | No |
| WikiPathways | https://www.wikipathways.org/ | No | No | Yes | No |
| SMPDB | https://smpdb.ca/ | No | No | Yes | No |
| LipidBank | https://lipidbank.jp/ | No | No | No | No |

| | | | | | |
|---|---|---|---|---|---|
| LipidMaps | https://www.lipidmaps.org/ | No | No | No | No |
| WikiPedia | https://www.wikipedia.org/ | No | No | No | No |
| GNPS | https://gnps.ucsd.edu/ProteoSAFe/static/gnps-splash.jsp?redirect=auth | No | Yes | No | No |
| MassBank | https://massbank.eu/MassBank/ | No | Yes | No | No |
| Mona | https://mona.fiehnlab.ucdavis.edu/ | No | Yes | No | No |
| Recon3 | https://www.vmh.life/ | No | No | No | Yes |
| MetaNetx | https://www.metanetx.org/ | No | No | No | Yes |
| Rhea | https://www.rhea-db.org/ | No | No | No | Yes |

**Supplementary Table 2. Comparison between TidyMass2 and existing tools.**

| Name | TidyMass2 | RforMassSpectrometry | MetaboAnalyst | MetOrigin2 | PIUMet | Mummichog | MSDIAL |
|---|---|---|---|---|---|---|---|
| Comprehensive workflow | +++ | ++ | +++ | + | + | + | ++ |
| Command-line interface | ✓ | ✓ | ✓ | - | - | ✓ | - |
| GUI/Online interface | ✓ | - | ✓ | ✓ | ✓ | ✓ | ✓ |
| Cross-platform utility | +++ | +++ | +++ | +++ | +++ | +++ | + |
| Local PC deployment | ✓ | ✓ | ✓ | - | - | ✓ | ✓ |
| Server deployment | ✓ | ✓ | ✓ | - | - | ✓ | - |
| Customized metabolite database | ✓ | - | - | - | - | - | ✓ |
| Customized pathway database | ✓ | - | - | - | - | - | - |
| Feature-based module analysis | ✓ | - | ✓ | - | ✓ | ✓ | - |
| Metabolite origin analysis | ✓ | - | - | ✓ | ✓ | - | - |
| Open-source (code) | +++ | +++ | +++ | - | - | +++ | +++ |
| Open-source (databases) | +++ | - | - | - | + | - | +++ |

**Note:** Symbols used for feature evaluations with "√" for present, "-" for absent, and "+" for a more quantitative assessment, with more "+" indicating better support.

**Supplementary Table 3. 27 dysregulated metabolic modules.**

| Name | P value | Metabolite number | Metabolite ID (KEGG) |
|---|---|---|---|
| metabolic_module_1 | 0 | 41 | C06099{}C00521{}C11398{}C11412{}C01123{}C01765{}C07271{}C11393{}C11399{}C11410{}C11411{}C11416{}C11384{}C00964{}C02452{}C09902{}C20261{}C02344{}C01766{}C09847{}C11408{}C11396{}C11395{}C11409{}C11397{}C11400{}C11413{}C11401{}C11418{}C11415{}C16462{}C16461{}C11924{}C01499{}C09848{}C02576{}C11383{}C01767{}C11338{}C11420{}C20260 |
| metabolic_module_2 | 0 | 84 | C00763{}C01905{}C00079{}C00041{}C00148{}C00152{}C00407{}C00135{}C00047{}C00065{}C00062{}C00123{}C00183{}C00078{}C00077{}C00792{}C01682{}C00064{}C00097{}C02486{}C00303{}C03793{}C05552{}C07541{}C06735{}C16438{}C16590{}C16593{}C03912{}C03564{}C19716{}C20277{}C20278{}C20313{}C17349{}C20978{}C22162{}C02356{}C00334{}C03406{}C15767{}C02155{}C03227{}C02794{}C01165{}C01157{}C00884{}C06121{}C02265{}C05933{}C04020{}C21015{}C11332{}C02993{}C16435{}C01015{}C05147{}C05636{}C07544{}C08290{}C16696{}C16831{}C17255{}C19706{}C19781{}C20330{}C20487{}C20488{}C20802{}C20850{}C21026{}C21092{}C21160{}C21283{}C04322{}C01924{}C02427{}C02728{}C01877{}C04148{}C02710{}C01570{}C03230{}C00977 |
| metabolic_module_3 | 0 | 28 | C00185{}C00031{}C00116{}C00267{}C00897{}C00208{}C01083{}C00103{}C01971{}C06421{}C02048{}C02074{}C02655{}C03855{}C08240{}C11546{}C15548{}C15970{}C00252{}C00221{}C01518{}C01594{}C08334{}C13857{}C04197{}C19636{}C02130{}C08325 |
| metabolic_module_4 | 0 | 109 | C00019{}C01917{}C00021{}C07349{}C02890{}C00580{}C00409{}C05204{}C06347{}C06629{}C07028{}C07966{}C03114{}C21584{}C09390{}C11839{}C11840{}C12044{}C12416{}C12476{}C05315{}C15682{}C15681{}C02483{}C20248{}C16230{}C16708{}C16713{}C16789{}C17702{}C17703{}C20444{}C20629{}C20673{}C20677{}C20819{}C21183{}C21198{}C21201{}C21281{}C21427{}C21428{}C21429{}C21433{}C21434{}C21436{}C21647{}C21648{}C22142{}C05298{}C00978{}C00189{}C00780{}C01161{}C16675{}C07481{}C13747{}C07130{}C07480{}C14152{}C00346{}C00588{}C13482{}C00385{}C02666{}C02918{}C01586{}C20674{}C02442{}C06212{}C09642{}C00758{}C00841{}C08608{}C21602{}C01957{}C05203{}C08299{}C05831{}C05902{}C01008{}C05594{}C08526{}C02079{}C02381{}C04208{}C04444{}C08263{}C10159{}C15530{}C15536{}C15616{}C15883{}C16419{}C19895{}C20234{}C20790{}C20872{}C01004{}C10652{}C06199{}C05317{}C06812{}C04293{}C08304{}C15531{}C10098{}C19807{}C03986 |
| metabolic_module_5 | 2.00E-05 | 77 | C00020{}C00314{}C00133{}C00740{}C00022{}C00515{}C02713{}C00246{}C02703{}C00109{}C00163{}C00466{}C03943{}C03341{}C03248{}C04137{}C00956{}C05556{}C09822{}C00164{}C00627{}C01089{}C1 |

| | | | |
|---|---|---|---|
| | | | 2477{}C12986{}C16138{}C16149{}C00441{}C02291{}C17237{}C19694{}C19719{}C02218{}C03283{}C05984{}C00542{}C02488{}C00250{}C01077{}C21096{}C01712{}C01607{}C01588{}C03401{}C08316{}C06429{}C08323{}C08367{}C16525{}C16536{}C03197{}C01118{}C01401{}C00534{}C08739{}C05714{}C16434{}C04210{}C02721{}C04076{}C03210{}C00736{}C00450{}C01275{}C16436{}C01136{}C01110{}C01092{}C00793{}C01037{}C03340{}C02132{}C02527{}C20258{}C20904{}C22087{}C03996{}C00716 |
| metabolic_module_6 | 1.00E-04 | 25 | C00794{}C00089{}C02209{}C00159{}C00095{}C00392{}C00668{}C01096{}C00085{}C00636{}C00092{}C01172{}C17209{}C04006{}C20236{}C20861{}C00275{}C00644{}C01742{}C03267{}C06468{}C06187{}C06019{}C19761{}C20887 |
| metabolic_module_7 | 0.00023 | 34 | C01732{}C00042{}C03325{}C01851{}C02046{}C06658{}C06659{}C06660{}C10842{}C11860{}C11857{}C11862{}C11859{}C11863{}C11866{}C11868{}C06094{}C15770{}C18312{}C11864{}C03648{}C11858{}C03761{}C00490{}C00859{}C06095{}C11865{}C02034{}C14162{}C02035{}C06096{}C11869{}C20632{}C11855 |
| metabolic_module_8 | 0.00046 | 21 | C01697{}C00243{}C01970{}C00764{}C00984{}C00124{}C05402{}C05401{}C01113{}C00962{}C01097{}C00795{}C01235{}C05399{}C05400{}C03384{}C01582{}C04452{}C21523{}C21524{}C02965 |
| metabolic_module_9 | 0.00053 | 33 | C00427{}C00219{}C11878{}C11881{}C18220{}C18222{}C06428{}C11887{}C14770{}C18211{}C18219{}C18221{}C11882{}C06425{}C16527{}C03242{}C14771{}C00909{}C05956{}C13856{}C14768{}C14769{}C14748{}C20388{}C18177{}C12083{}C14822{}C12077{}C09118{}C20329{}C14807{}C14778{}C14749 |
| metabolic_module_10 | 0.00077 | 20 | C00270{}C00140{}C03878{}C00329{}C08349{}C04501{}C00357{}C01674{}C00611{}C19910{}C19909{}C04017{}C04015{}C04256{}C00645{}C06372{}C04257{}C01075{}C04016{}C04886 |
| metabolic_module_11 | 0.00224 | 35 | C00712{}C00487{}C15025{}C00318{}C02290{}C00823{}C04114{}C01181{}C06424{}C06427{}C21761{}C00249{}C01530{}C08320{}C06426{}C16300{}C16522{}C02862{}C19670{}C06123{}C00517{}C05828{}C01149{}C01259{}C01948{}C08388{}C02571{}C19936{}C20704{}C22135{}C20826{}C02838{}C03017{}C03195{}C17535 |
| metabolic_module_12 | 0.00389 | 17 | C15788{}C15787{}C15789{}C15790{}C15791{}C15799{}C15798{}C15797{}C15800{}C16253{}C15792{}C16251{}C15801{}C15793{}C17733{}C15786{}C17735 |
| metabolic_module_13 | 0.00924 | 23 | C06069{}C01500{}C06070{}C11389{}C11404{}C11402{}C17622{}C20222{}C20223{}C10469{}C10453{}C17621{}C03985{}C04433{}C09861{}C01433{}C06071{}C10428{}C09987{}C20221{}C20225{}C10478{}C09910 |
| metabolic_module_14 | 0.01093 | 12 | C14786{}C04314{}C00829{}C14787{}C11714{}C11713{}C06205{}C14791{}C14793{}C14792{}C04514{}C14784 |
| metabolic_module | 0.01174 | 10 | C21030{}C01188{}C06002{}C18318{}C02170{}C06001{}C03284{}C01205{}C00349{}C20846 |

| | | | |
|---|---|---|---|
| _15 | | | |
| metabolic_module_16 | 0.01175 | 28 | C00881{}C01121{}C11992{}C03240{}C06630{}C06616{}C00363{}C11955{}C11963{}C11972{}C11991{}C11993{}C11997{}C12000{}C12002{}C12384{}C12385{}C12404{}C12407{}C12412{}C12424{}C12472{}C12481{}C18034{}C18635{}C18634{}C06635{}C21350 |
| metabolic_module_17 | 0.01764 | 14 | C00280{}C00674{}C00523{}C07632{}C01227{}C03772{}C04295{}C05139{}C05290{}C05140{}C20144{}C11135{}C20252{}C18045 |
| metabolic_module_18 | 0.02119 | 11 | C19756{}C19743{}C01126{}C19746{}C09704{}C03220{}C20325{}C16537{}C03461{}C19691{}C20121 |
| metabolic_module_19 | 0.02135 | 11 | C03375{}C00750{}C00986{}C02567{}C01029{}C00612{}C05665{}C03413{}C05936{}C02946{}C18170 |
| metabolic_module_20 | 0.02455 | 8 | C00546{}C00583{}C00424{}C05235{}C00937{}C02912{}C02917{}C15499 |
| metabolic_module_21 | 0.02522 | 13 | C05487{}C00735{}C18075{}C05488{}C05138{}C02373{}C05501{}C05499{}C05284{}C02821{}C14256{}C05489{}C02822 |
| metabolic_module_22 | 0.02527 | 24 | C00132{}C01761{}C02151{}C03677{}C05616{}C09694{}C11632{}C11633{}C11807{}C16502{}C16503{}C16504{}C16505{}C18640{}C06800{}C19613{}C11045{}C13244{}C03686{}C17530{}C16508{}C16506{}C16507{}C19681 |
| metabolic_module_23 | 0.03215 | 19 | C00235{}C06067{}C09023{}C20512{}C20532{}C20543{}C20545{}C20563{}C20982{}C20983{}C18083{}C04290{}C02008{}C06068{}C18053{}C19767{}C20594{}C20636{}C16521 |
| metabolic_module_24 | 0.0364 | 26 | C01054{}C08624{}C11505{}C01902{}C01724{}C11455{}C08628{}C17966{}C19820{}C19832{}C19833{}C20187{}C20188{}C20189{}C20191{}C20192{}C20193{}C20194{}C20195{}C20200{}C19835{}C08637{}C19918{}C19801{}C08615{}C08626 |
| metabolic_module_25 | 0.03701 | 6 | C00101{}C00143{}C21801{}C21802{}C00664{}C00445 |
| metabolic_module_26 | 0.03713 | 18 | C08062{}C00108{}C00196{}C00885{}C00251{}C00805{}C10733{}C00134{}C20579{}C00254{}C21147{}C03005{}C00555{}C15668{}C00826{}C10497{}C01405{}C03002 |
| metabolic_module_27 | 0.04192 | 10 | C02991{}C01934{}C19758{}C02338{}C02476{}C00507{}C02431{}C20354{}C03979{}C00861 |

**Supplementary Table 4. Identified pathways for dysregulated metabolic modules.**

| Pathway ID | Pathway name | p_value_adjust | Metabolite ID (KEGG) |
| --- | --- | --- | --- |
| hsa00330 | Arginine and proline metabolism | 1.97E-11 | C03912;C00986;C03564;C15767;C02946;C00334;C00555;C01877;C01110;C04137;C00763;C10497;C00884;C01157;C00062;C00441;C01165;C00077;C00148;C05933;C05936;C01682;C03375;C00134;C00022;C00019;C00750;C19706;C05147 |
| hsa00052 | Galactose metabolism | 1.73E-10 | C05401;C00095;C00085;C00124;C01113;C00031;C00103;C00159;C00794;C00795;C01097;C05400;C01697;C00116;C00243;C05399;C05402;C00089;C00984;C01235;C00267;C00668 |
| hsa05230 | Central carbon metabolism in cancer | 9.85E-09 | C00085;C00031;C00092;C00041;C00062;C00152;C00097;C00064;C00135;C00407;C00123;C00079;C00148;C00065;C00078;C00183;C00022;C00042 |
| hsa00470 | D-Amino acid metabolism | 1.49E-08 | C03943;C03564;C03341;C01110;C00133;C00792;C00793;C00515;C02265;C00763;C00740;C01157;C00041;C00062;C00097;C00064;C00135;C00047;C00077;C00079;C00148;C00065;C00134;C00022 |
| hsa02010 | ABC transporters | 2.30E-07 | C04114;C01181;C00487;C00185;C01674;C00095;C00031;C00159;C04137;C00794;C00881;C00116;C01157;C00041;C00062;C00064;C00135;C00407;C00123;C00047;C00077;C00079;C00148;C00065;C00183;C00243;C00208;C00392;C05402;C00140;C01682;C00134;C00089;C01083 |
| hsa04974 | Protein digestion and absorption | 5.83E-07 | C00246;C00041;C00062;C00152;C00097;C00064;C00135;C00407;C00123;C00047;C00079;C00148;C16138;C00065;C00078;C00183;C00163;C00134 |
| hsa00051 | Fructose and mannose metabolism | 3.48E-05 | C00424;C03979;C00095;C00644;C00159;C00636;C00275;C00794;C02431;C01934;C02991;C00507;C00861;C00392;C01096;C00267;C03267 |
| hsa04978 | Mineral absorption | 4.85E-05 | C00124;C00031;C00041;C00152;C00064;C00407;C00123;C00079;C00148;C00065;C00078;C00183 |

| | | | |
|---|---|---|---|
| hsa00970 | Aminoacyl-tRNA biosynthesis | 8.09E-05 | C00041;C00062;C00152;C00097;C00064;C00135;C00407;C00123;C00047;C00079;C00148;C16138;C00065;C00078;C00183;C00101 |
| hsa00140 | Steroid hormone biosynthesis | 0.00040139 | C05488;C05489;C18075;C05284;C05140;C05139;C05499;C05487;C05138;C05290;C05298;C05501;C02373;C00674;C03772;C18045;C04295;C00280;C00523;C11135;C00735;C01227 |
| hsa04742 | Taste transduction | 0.00111382 | C00334;C00020;C11045;C00133;C00095;C00031;C02265;C00740;C00208;C00780;C00089 |
| hsa00270 | Cysteine and methionine metabolism | 0.00138186 | C02356;C00109;C00793;C02218;C00041;C00441;C02291;C00097;C00065;C00409;C01077;C01118;C00022;C00021;C00019;C21015 |
| hsa00260 | Glycine, serine and threonine metabolism | 0.00222167 | C00986;C00109;C00143;C00740;C03283;C00441;C02291;C00097;C00065;C00078;C00546;C00022;C00101 |
| hsa04973 | Carbohydrate digestion and absorption | 0.00443079 | C00246;C00095;C00124;C00031;C00092;C00243;C00208;C00163;C00089 |
| hsa01040 | Biosynthesis of unsaturated fatty acids | 0.00570773 | C08323;C06429;C06428;C06426;C00712;C06427;C16527;C00219;C03242;C08316;C00249;C16525;C06425;C16522;C01530;C08320 |
| hsa04913 | Ovarian steroidogenesis | 0.00876224 | C14770;C05138;C14768;C14769;C04295;C00280;C00219;C01227 |
| hsa00310 | Lysine degradation | 0.01042917 | C01259;C00450;C01149;C01181;C00164;C00487;C04020;C00956;C04076;C00047;C03793;C00042 |

| | | | |
|---|---|---|---|
| hsa00500 | Starch and sucrose metabolism | 0.01042917 | C00185;C00095;C00085;C00031;C00103;C00092;C00252;C00208;C00089;C01083 |
| hsa00640 | Propanoate metabolism | 0.01820643 | C00424;C06002;C05984;C00109;C05235;C00546;C02170;C00583;C00163;C00042 |
| hsa04270 | Vascular smooth muscle contraction | 0.01820643 | C14770;C14771;C14748;C14768;C14769;C00219 |
| hsa00250 | Alanine, aspartate and glutamate metabolism | 0.02095442 | C03912;C00334;C00041;C00152;C00064;C03406;C00022;C00042 |
| hsa04726 | Serotonergic synapse | 0.02363209 | C14770;C14771;C14768;C14769;C00219;C00078;C00909;C05956;C00427;C00780 |
| hsa00280 | Valine, leucine and isoleucine degradation | 0.02363209 | C01205;C06001;C06002;C00349;C00164;C03284;C00407;C00123;C00183;C02170 |
| hsa04150 | mTOR signaling pathway | 0.02571935 | C00020;C00062;C00123 |
| hsa00010 | Glycolysis / Gluconeogenesis | 0.03577246 | C06187;C00031;C00103;C00022;C00267;C00668;C00221;C01172 |
| hsa04723 | Retrograde endocannabinoid signaling | 0.03807027 | C13856;C00334;C00219;C00189;C00116;C00427 |

| | | | |
|---|---|---|---|
| hsa04922 | Glucagon signaling pathway | 0.04461446 | C00085;C00031;C00103;C00022;C00042;C00668;C01172 |
| hsa00670 | One carbon pool by folate | 0.04461446 | C00445;C00143;C00664;C00101 |
| hsa04931 | Insulin resistance | 0.04538455 | C00095;C00085;C00031;C00092;C02571;C00022 |

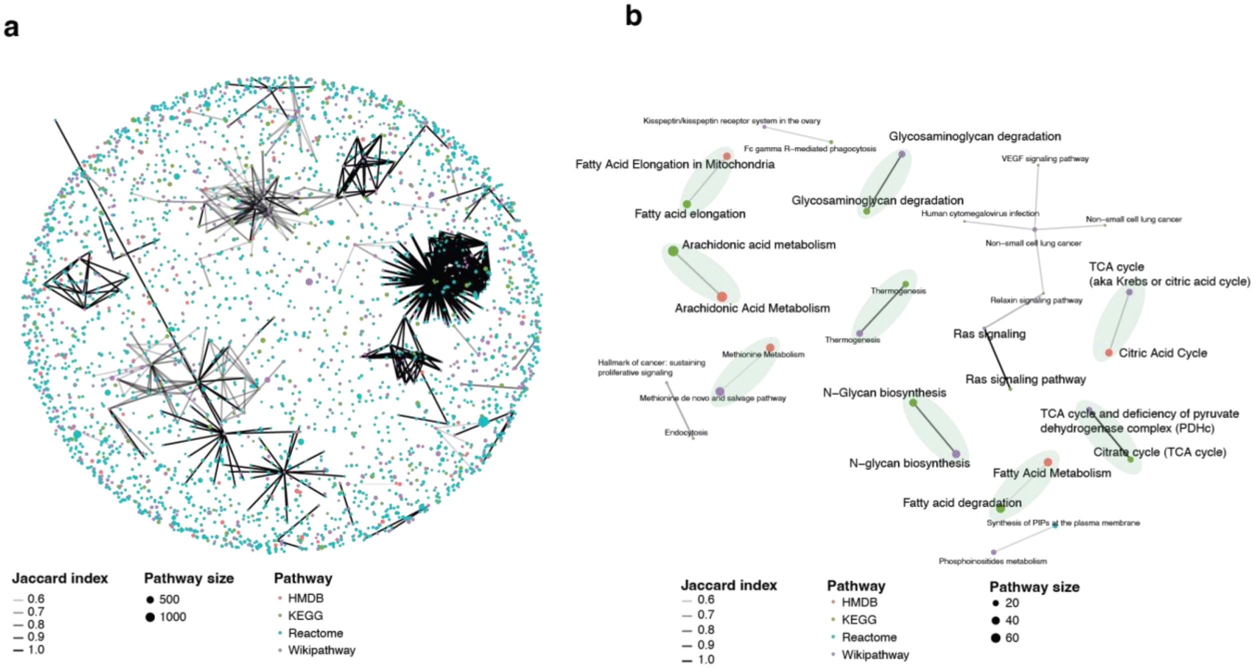


**Supplementary Figure 1 | Similarity network of metabolic pathways across four public databases. a,** Global network visualization of metabolic pathways from HMDB, KEGG, Reactome, and WikiPathways. Nodes represent individual pathways, colored by their database of origin. Edges denote pairwise similarity based on the Jaccard index, with edge thickness indicating the degree of overlap. Node size reflects the number of metabolites within each pathway. **b,** Enlarged subnetwork highlighting clusters of highly similar or identical pathways shared across multiple databases. These clusters illustrate redundancy and overlap in pathway definitions, underscoring the need for integrated pathway analysis in untargeted metabolomics.

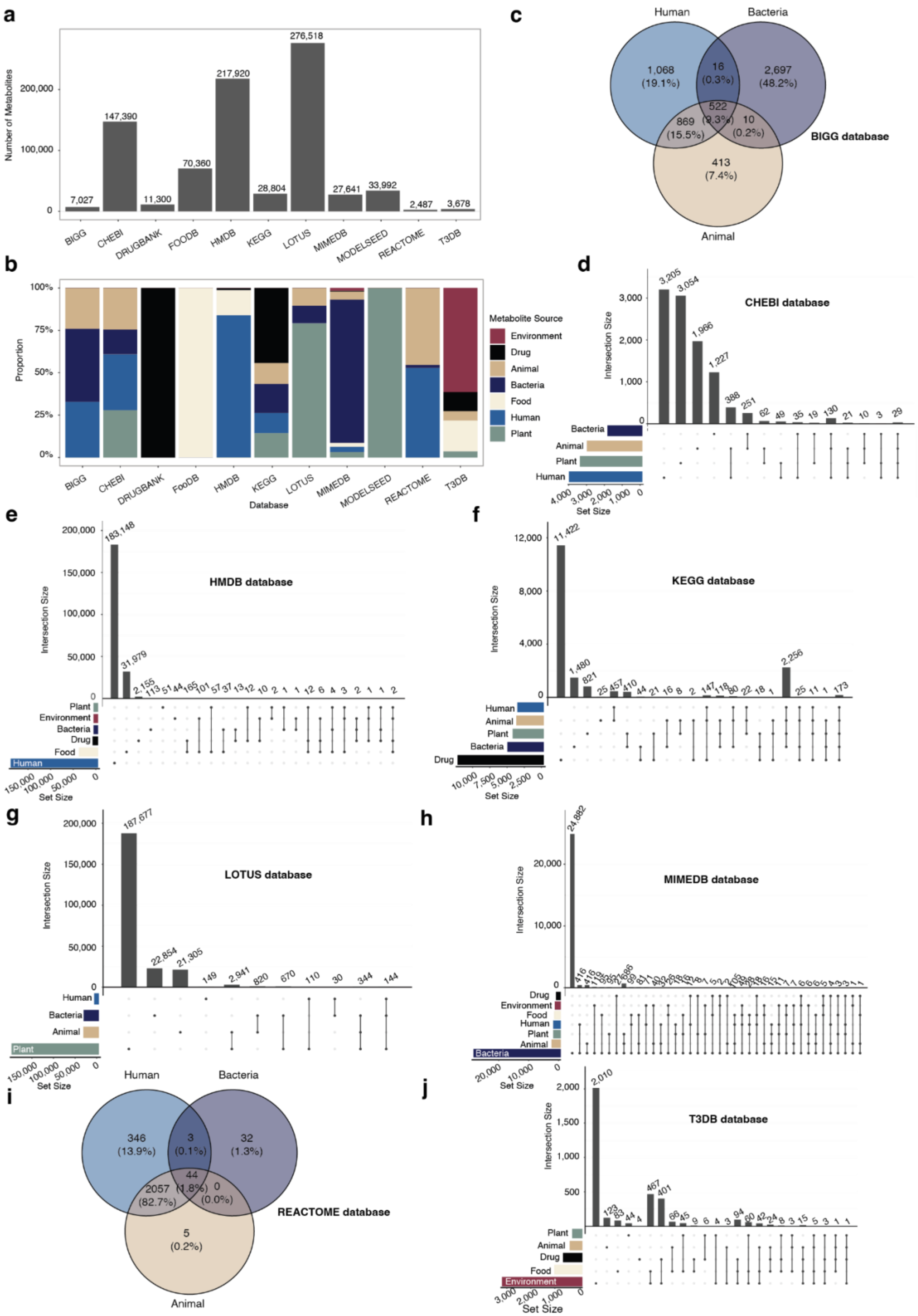

a
Number of Metabolites
200,000
100,000
0
7,027
147,390
11,300
70,360
217,920
28,804
276,518
27,641
33,992
2,487
3,678
BIGG
CHEBI
DRUGBANK
FOODB
HMDB
KEGG
LOTUS
MIMEDB
MODELSEED
REACTOME
T3DB
b
Proportion
100%
75%
50%
25%
0%
Database
Metabolite Source
Environment
Drug
Animal
Bacteria
Food
Human
Plant
c
Human
Bacteria
Animal
1,068
(19.1%)
16
(0.3%)
2,697
(48.2%)
522
(9.3%)
869
(15.5%)
10
(0.2%)
413
(7.4%)
BIGG database
d
CHEBI database
Intersection Size
Set Size
3,205
3,054
1,966
1,227
388
251
62
49
35
19
130
21
10
3
29
e
HMDB database
Intersection Size
Set Size
183,148
31,979
f
KEGG database
Intersection Size
Set Size
11,422
1,480
821
2,256
g
LOTUS database
Intersection Size
Set Size
187,577
22,854
21,305
149
2,941
820
670
110
30
344
144
h
MIMEDB database
Intersection Size
Set Size
24,882
i
Human
Bacteria
Animal
346
(13.9%)
3
(0.1%)
32
(1.3%)
44
(1.8%)
2057
(82.7%)
0
(0.0%)
5
(0.2%)
REACTOME database
j
T3DB database
Intersection Size
Set Size
2,010
467
401

**Supplementary Figure 2 | Detailed overview of the integrated metabolite database with source information.** **a,** Number of metabolites contributed by each of the 11 integrated public databases. **b,** Proportional distribution of source categories (human, animal, bacteria, plant, food, drug, and environment) within each database. **c-j,** Source overlap analysis across individual databases. **c,** The Venn diagram shows the source category overlap in the BiGG database. **d-f,** UpSet plots illustrating source intersections for metabolites in the ChEBI **(d)**, HMDB **(e)**, and KEGG **(f)** databases. **g-h,** UpSet plots showing source distribution in the LOTUS **(g)** and MiMeDB **(h)** databases. **i,** Venn diagram of source category overlap for the Reactome database. **j,** UpSet plot for source classification in the T3DB database.

**a**

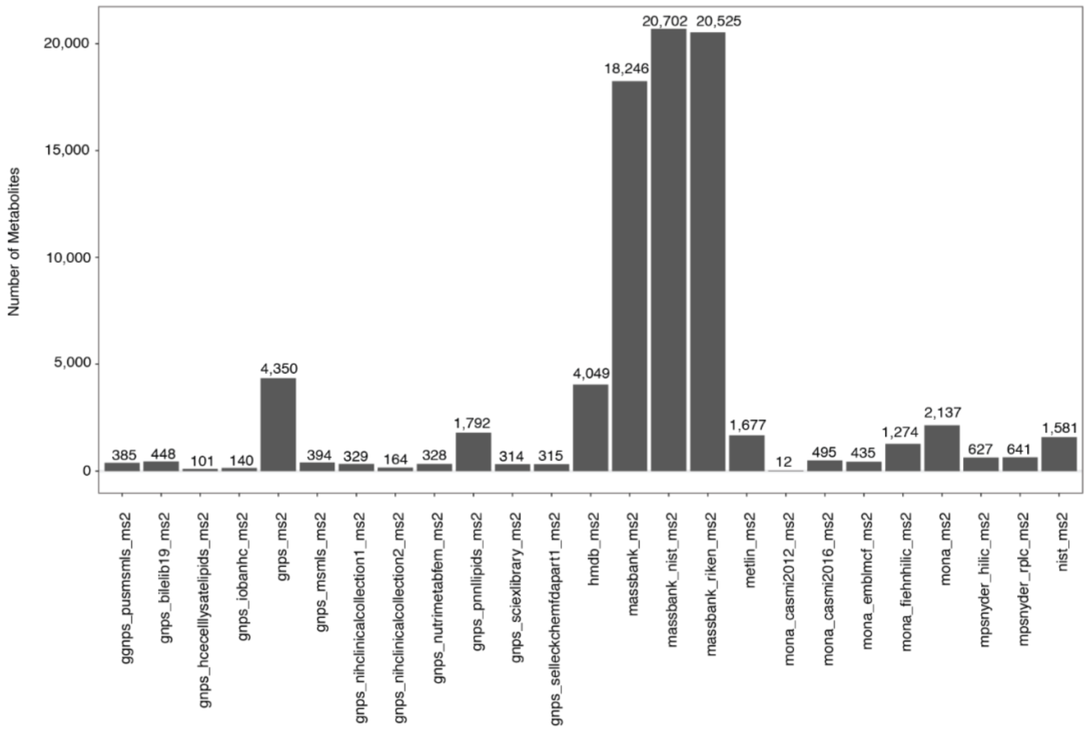

Number of Metabolites
0
5,000
10,000
15,000
20,000
385
448
101
140
4,350
394
329
164
328
1,792
314
315
4,049
18,246
20,702
20,525
1,677
12
495
435
1,274
2,137
627
641
1,581
ggnps_pusmsmls_ms2
gnps_bileilb19_ms2
gnps_hcecelllysatelipids_ms2
gnps_iobanhc_ms2
gnps_ms2
gnps_msmls_ms2
gnps_nihclinicalcollection1_ms2
gnps_nihclinicalcollection2_ms2
gnps_nutrimetabfem_ms2
gnps_pnnllipids_ms2
gnps_sciexlibrary_ms2
gnps_selleckchemfdapart1_ms2
hmdb_ms2
massbank_ms2
massbank_nist_ms2
massbank_riken_ms2
metlin_ms2
mona_casmi2012_ms2
mona_casmi2016_ms2
mona_emblmcf_ms2
mona_fiehnhilic_ms2
mona_ms2
mpsnyder_hilic_ms2
mpsnyder_rplc_ms2
nist_ms2


**b**

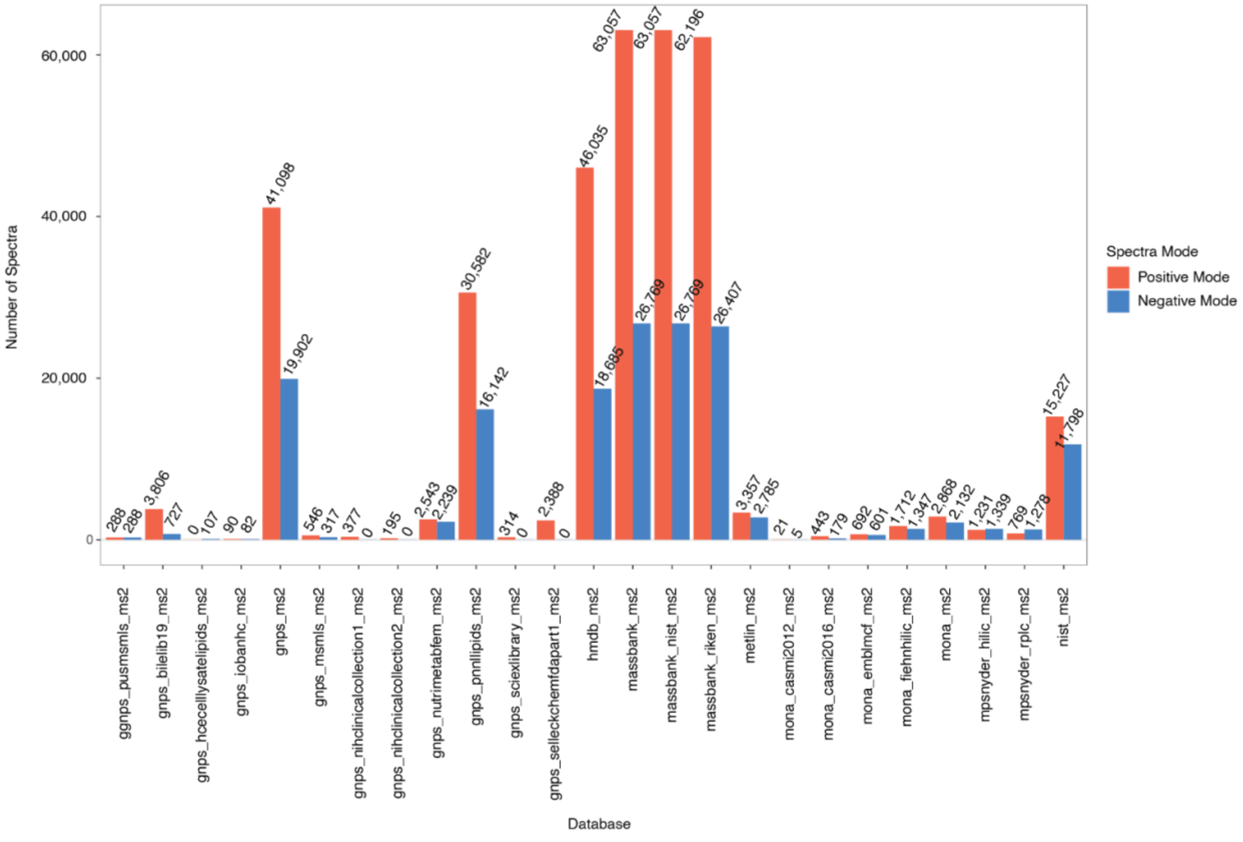

Number of Spectra
0
20,000
40,000
60,000
288
288
3,806
727
0
107
90
82
41,098
19,902
546
317
377
0
195
0
2,543
2,239
30,582
16,142
314
0
2,388
0
46,035
18,685
63,057
26,769
63,057
26,769
62,196
26,407
3,357
2,785
21
5
443
179
692
601
1,712
1,347
2,868
2,132
1,231
1,339
769
1,278
15,227
11,798
Spectra Mode
Positive Mode
Negative Mode
ggnps_pusmsmls_ms2
gnps_bileilb19_ms2
gnps_hcecelllysatelipids_ms2
gnps_iobanhc_ms2
gnps_ms2
gnps_msmls_ms2
gnps_nihclinicalcollection1_ms2
gnps_nihclinicalcollection2_ms2
gnps_nutrimetabfem_ms2
gnps_pnnllipids_ms2
gnps_sciexlibrary_ms2
gnps_selleckchemfdapart1_ms2
hmdb_ms2
massbank_ms2
massbank_nist_ms2
massbank_riken_ms2
metlin_ms2
mona_casmi2012_ms2
mona_casmi2016_ms2
mona_emblmcf_ms2
mona_fiehnhilic_ms2
mona_ms2
mpsnyder_hilic_ms2
mpsnyder_rplc_ms2
nist_ms2
Database

**Supplementary Figure 3 | Overview of the merged $MS^2$ spectral database integrated in TidyMass2.**
**a,** Total number of unique metabolites represented across 25 public and in-house $MS^2$ spectral databases.
**b,** Distribution of $MS^2$ spectra across positive and negative ionization modes for each database.

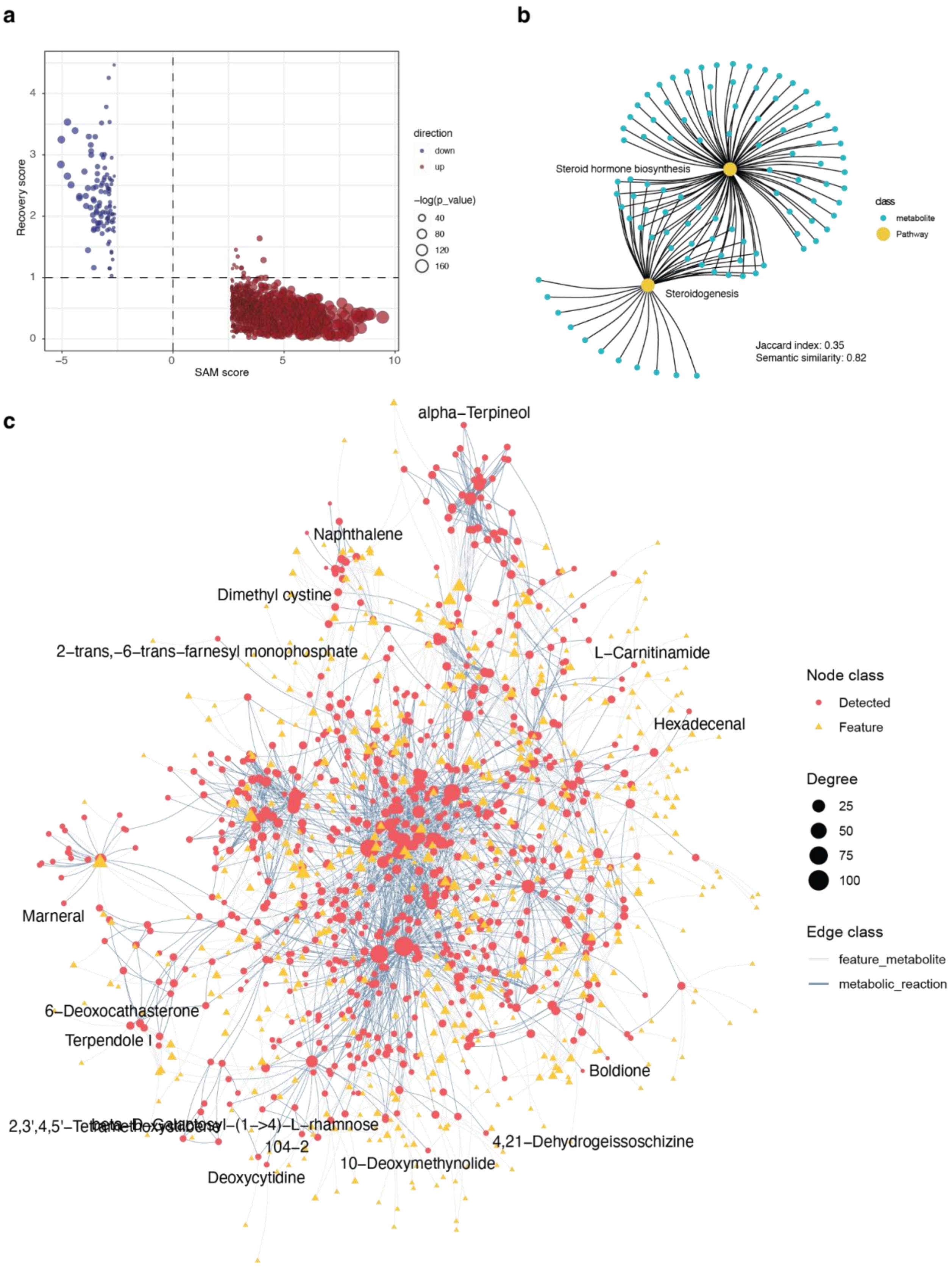

a
direction
down
up
−log(p_value)
40
80
120
160
Recovery score
SAM score
b
Steroid hormone biosynthesis
Steroidogenesis
class
metabolite
Pathway
Jaccard index: 0.35
Semantic similarity: 0.82
c
alpha−Terpineol
Naphthalene
Dimethyl cystine
2−trans,−6−trans−farnesyl monophosphate
L−Carnitinamide
Hexadecenal
Marneral
6−Deoxocathasterone
Terpendole I
Boldione
104−2
4,21−Dehydrogeissoschizine
10−Deoxymethynolide
Deoxycytidine
Node class
Detected
Feature
Degree
25
50
75
100
Edge class
feature_metabolite
metabolic_reaction

**Supplementary Figure 4 | Dysregulated metabolic features and network remodeling during pregnancy. a,** Scatter plot showing SAM scores and recovery scores for all significantly dysregulated metabolic features across pregnancy. A positive SAM score indicates increased abundance during pregnancy, while a negative SAM score indicates decreased levels. Recovery scores reflect post-delivery changes relative to late pregnancy: scores < 0 suggest postpartum decline, and scores >0 indicate postpartum elevation. **b,** Network visualization showing the overlap between two pathways from different databases, demonstrating pathway convergence across sources. **c,** Global dysregulated metabolic network identified using the metabolic feature-based module analysis in TidyMass2. Nodes represent either metabolic features (yellow triangles) or putatively annotated metabolites (red circles), with node size reflecting connectivity (degree). Edges denote biological relationships based on reaction pairs or metabolite-feature annotations.

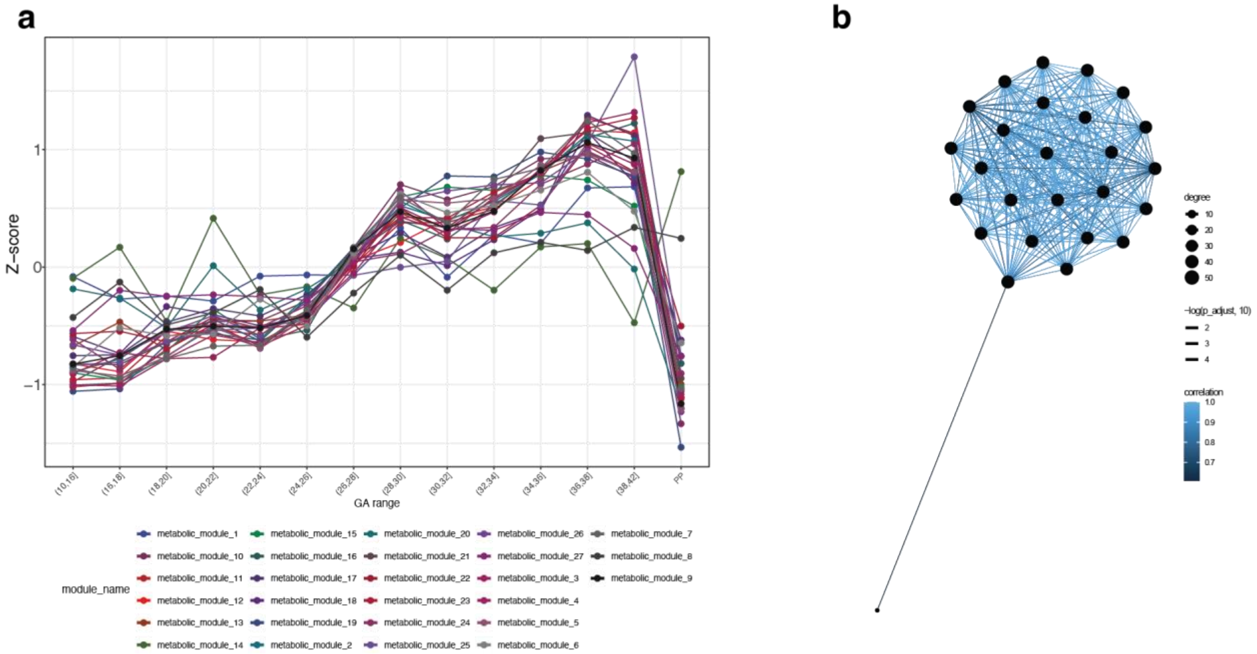


**Supplementary Figure 5 | Temporal dynamics and interconnectivity of dysregulated metabolic modules during pregnancy. a,** Line plot showing longitudinal changes in the abundance of 27 dysregulated metabolic modules across gestation. Each line represents the Z-score trajectory of a single module across gestational age (GA) intervals, with a sharp decline observed postpartum (PP). **b,** Correlation network illustrating relationships among the 27 metabolic modules. Node size indicates degree (number of connections), edge width reflects the significance of correlations (adjusted p-value), and edge color denotes the strength of correlations. This network highlights coordinated metabolic remodeling during pregnancy.
